# Curvature-mediated prewetting organize mitochondrial nucleoid

**DOI:** 10.64898/2026.04.06.716811

**Authors:** Xiangzi Hu, Li Shu, Guicong Zhang, Yongqian Jiang, Yuanfeng Yin, Yanyan Xu, Yong Wang, Yuxing Shang, Jiayi Cao, Ting Li, Sanhua Fang, Shuo Guo, Di Li, Dechen Jiang, Christoph A. Weber, Cong Liu, Zhongwen Chen, Xueping Zhao, Yifan Ge

## Abstract

The emergence of spatiotemporal order in diffusive cellular environments requires physical mechanisms to overcome the entropic drive toward disorder. While the organizing role of chemical signaling is well characterized, how complex membrane curvature regulate soluble molecular organization remains a fundamental open question. Here, we establish membrane curvature as a thermodynamic control parameter for the condensation of soluble proteins through prewetting. Using the mitochondrial transcription factor TFAM, we demonstrate that intrinsic membrane curvature locally drives a prewetting transitiona surface-mediated phase separation distinct from bulk condensation. By combining in vitro reconstitution, live-cell super-resolution imaging, cryo-electron tomography, and thermodynamic theory, we show that negatively high-curvature regions lower the nucleation energy barrier of Tfam, driving localized protein condensation at physiological concentrations. These results reveal that membrane curvature does not merely scaffold cellular machinery but manipulates the local free energy landscape, establishing a geometric principle for the spatial control of biological processes.

## Introduction

Living cells orchestrate complex biochemical reactions with remarkable spatial precision[1] despite operating in a crowded and highly dynamic cytoplasmic environment [2, 3].While liquid–liquid phase separation (LLPS) has emerged as a general physical mechanism driving functional subcellular organization [4–6], classical 3D bulk LLPS is fundamentally a spatial unrestricted process relies heavily on global concentration thresholds[5]. Consequently, it remains largely unclear how such free-floating condensates are precisely localized and robustly stabilized within complex subcellular architectures. Additionally, this framework faces a thermodynamic paradox: many functional assemblies form at concentrations far below the saturation level predicted by bulk phase diagrams[7]. Moreover, in organelles that undergoing constant fusion and fission, such as mitochondria[8],such membrane dynamics events induce drastic macroscopic fluctuations in volume and bulk concentrations, alongside strong hydrodynamic perturbation. Strikingly, despite these dramatic morphological changes,condensates such as mitochondria nucleoids, a structure critical to mitochondrial genomic stablity and function, exhibits remarkable structure fidelity and compositional integrity[9, 10]. If nucleoid formation solely relied on classical 3D bulk phase separation, such dynamic environments would inevitably lead to droplet dissolution and remarkable spatial drift.

To resolve these physical constraints, recent advances in soft matter physics propose that lipid membranes can serve as active platform, driving biomolecular assembly through a *prewetting* transition[11]. In this regime, interfacial interactions drastically compensate for the bulk chemical potential deficit, enabling surface condensation even when the bulk fluid is highly undersaturated [12–14]. Although this framework establishes membranes as general organizational cues, it remains largely unexplored how specific morphological features within the dynamic, fluidic landscape of the lipid bilayer actively regulate and thermodynamically dictate the spatial positioning of the surface-associated condensates.

Addressing such critical gap requires deciphering the exact coupling between the morphology of membrane-bound subcellular architecture and the functional protein phase behavior[15]. While the ability of biomolecular condensates to remodel membrane shape via capillary forces is well-documented[15–17], the reciprocal effect, how membrane curvature dictates the phase behavior of soluble proteins, still lacks a mechanistic description. To investigate this reciprocal coupling, we focus on the mitochondrion, which provides an ideal paradigm defined by the intricate, highly curved architecture of its inner membrane[18]. In the mitochondrial matrix, transcription Factor A (Tfam) a major nucleoid protein, compacts the mitochondrial genome into nucleoids through liquid–liquid phase separation [19, 20], supporting mitochondrial metabolic homeostasis[21]. However, how these soluble assemblies maintain stability *in vivo* amid the constant transport, fusion, and fission of the mitochondrial network remains difficult to reconcile with conventional solution chemistry.

In this study, we identify membrane curvature as a critical control parameter for biomolecular condensation. By employing in vitro reconstituted membranes with engineered curvature and live-cell super-resolution imaging, we demonstrate that Tfam, although without membrane association structures, undergoes a curvature-dependent prewetting transition. We find that membrane topography does not merely recruit proteins but alters the local phase diagram, allowing condensation to occur at physiologically low concentrations specifically at curved regions (**Fig. 1**). These results establish a mechanism where cellular geometry acts as a thermodynamic regulator, enabling precise spatial control over biochemical organization through the physics of wetting.

**Fig. 1.**
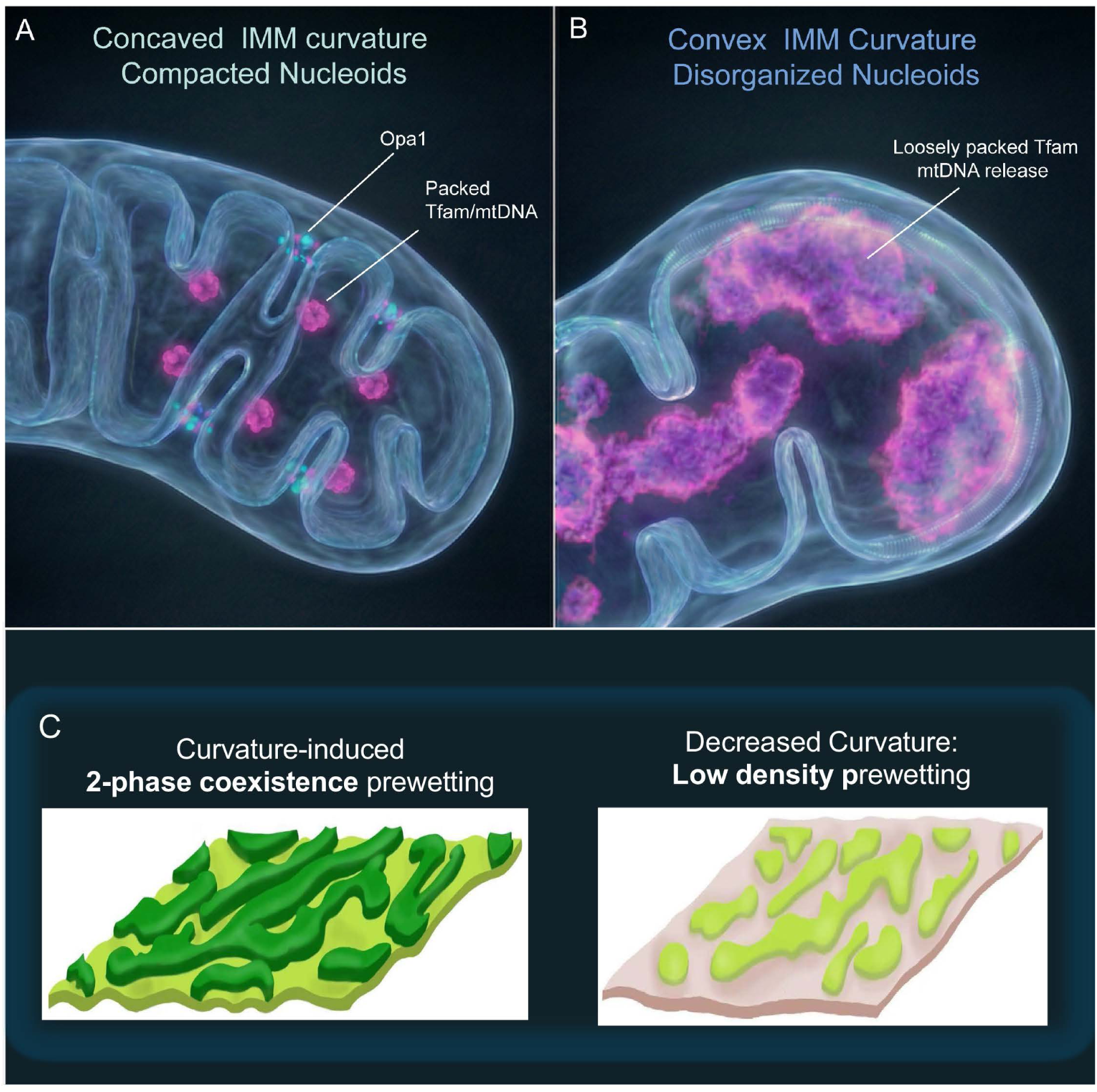
Curvature-dependent Tfam prewetting as the main driving force for nucleoids assembly. **A.** Illustration shows that normal IMM structures with highly curved cristae with well-packed nucleoids. While enlarged nucleoids are often observed in accompany with IMM disorganization as shown in **B**. **C.** Mechanistic schematic of two-phase surface condensation through prewetting on curved membrane surfaces. High membrane curvature (left) stabilizes a two-phase coexistence regime, promoting heterogeneous high-density prewetted surface condensates. Conversely, decreased curvature (right) suppresses this membrane-protein coupling, yielding a heterogeneous, low-density prewetted state.

## Results

### Membrane Curvature Regulates Mitochondrial Nucleoid Assembly via Surface-Mediated Phase Separation

Chemical cues are traditionally considered the primary triggers for catalytic phase separation [4, 22], membrane surfaces and relatively transient membrane-local cues like calcium concentration [23], receptor heterogeneity [13, 14, 24], and curvature[25] can act as potent physical catalyst to drive localized condensation well below bulk saturation levels. In mitochondria, nucleoids are LLPS-derived concensation initiated by Tfam [10, 20], and positioned in close association with the inner mitochondrial membrane(IMM)[26, 27] where nucleoid stability is sensitive to biochemical pathways that remodel IMM architecture[28], including the IMM fusion GTPase Opa1[29],whose loss reduced nucleoid abundance and promoted mtDNA release[29]. Motivated by these observations, we use Opa1 as a genetically and biochemically tractable handle to remodel IMM with the hypothesis that Tfam condensation acts in response to Opa1-mediated IMM structural regulation.

Using Structured Illumination Microscopy (SIM) on mTEC cells, we observed that while control cells exhibit packed nucleoids associated with IMM **(Fig. 2A)**, Opa1 knockout leads to mtDNA releasing and a homogeneous Tfam distribution within the mitochondria **(Fig. 2 A.B, Extended Fig. 1A)**. Reintroducing Opa1 isoform 1 variants (s-Opa1 and l-Opa1) into KO cells (**Extended Fig. 1B** ) restored Tfam nucleoid-like organization **(Fig. 2A,B)**. Notably, s-Opa1 rescued Tfam assembly despite failing to restore the mitochondrial membrane network **(Fig. 2A)**. Since s-Opa1 assemblies in the intermembrane space are physically separated from matrix-localized Tfam[30], our results suggests regulation occurs via membrane curvature rather than direct chemical interaction.

**Fig. 2.**
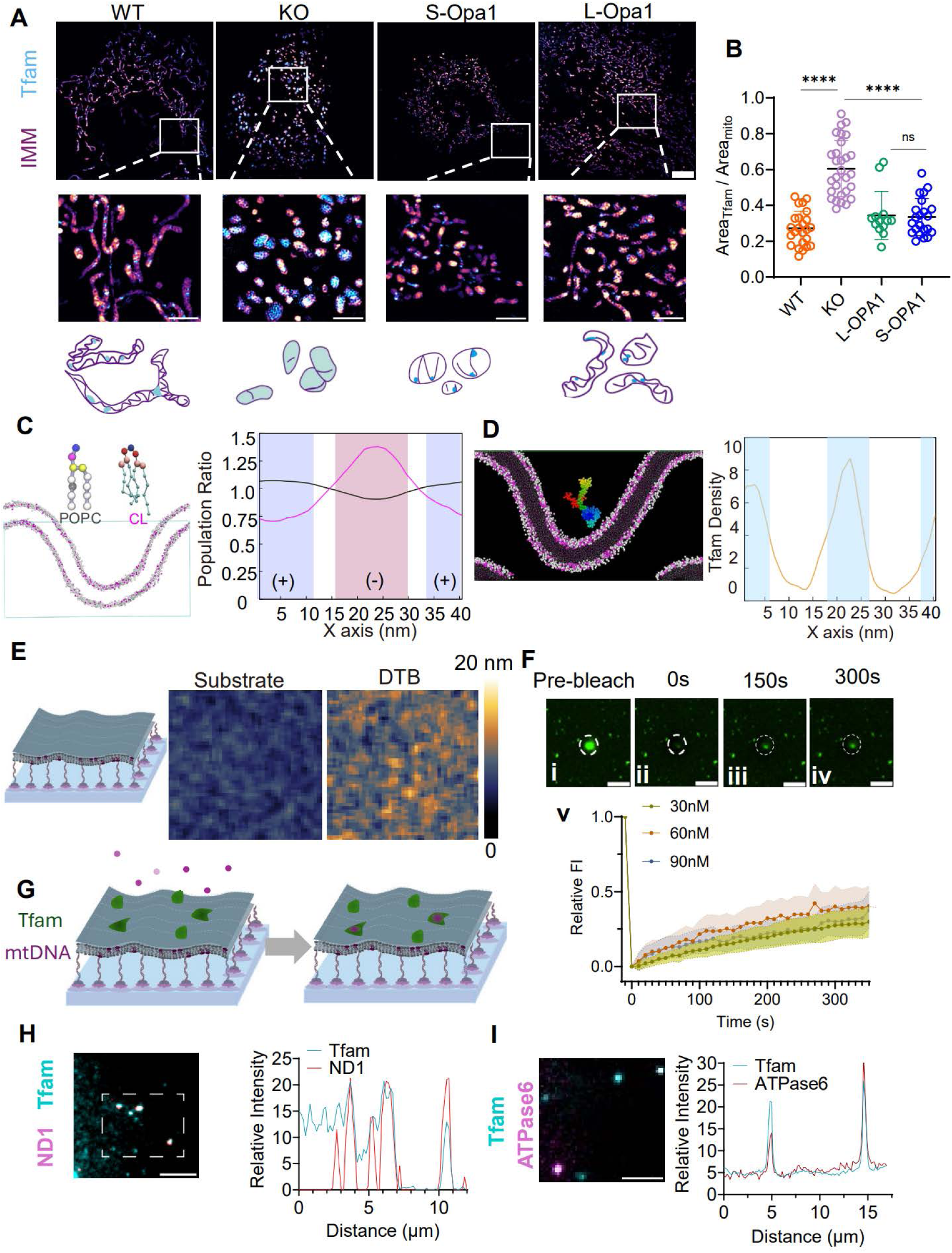
Tfam organization is correlated to IMM structure, possibly regulated by membrane surface condensation. **(A)** Live cell SIM images of cells labeled for Tfam (cyan) and the IMM (magenta). While WT cells show discrete Tfam puncta, Opa1 KO cells display diffuse Tfam distribution. Top row scale bars: 5 *µ*m; Middle row scale bars: 2 *µ*m. Bottom, schemes of IMM geometry and Tfam distribution with different Opa1 expression. **(B)** Quantification of Tfam matrix occupancy (Area_Tfam_/Area_mito_). **(C)** Molecular dynamics (MD) simulations of a POPC/CL bilayer show CL is enriched in negatively curved regions of each leaflet. Right: Quantification of lipid distribution in the upper leaflet; regions of negative and positive curvature are marked as (-) and (+), respectively. **(D)** MD snapshots showing preferential contact and conformational flexibility of Tfam at curved membrane regions. Right: Contact probability density profile. **(E)** Characterization of DNA-tethered Lipid Bilayers (DLBs) as IMM mimetics. SICM topology maps reveal that while tether-only substrates are flat, the resulting bilayers exhibit DNA-induced curvature (roughness), (FOV:1.35×1.35 *µm*^2^). **(F)** FRAP analysis of Tfam condensates on DLBs (30-90 nM). The partial fluorescence recovery indicates liquid-like dynamics (Scale bar:1 *µm*). **(G)** Schematic of mtDNA recruitment by surface-condensed Tfam. **H, I** Validation of mtDNA recruitment via colocalization of Tfam with ND1 (H) and ATPase6 (I) fractions.(B: One-way ANOVA; **** *P <* 0.0001, ns: not significant. Each data p5resents a biological replicates.)

To test this hypothesis, we first performed coarse-grained molecular dynamics (MD) simulations using IMM mimetics composed of POPC and 20 mol% cardiolipin (CL), a lipid uniquely enriched in the IMM. Our simulations recapitulated the curvature sorting of CL[31], showing enrichment at negatively curved segments of each leaflet **(Fig. 2C)**. Introducing a single Tfam molecule[32] revealed pronounced conformational flexibility and random surface contacts. Interestingly, unlike other curvature-sensitive proteins, Tfam does not physically adsorb to the membrane surface but contacts it transiently **(Extended Movies 1 and 2)**. Quantification of contact probability showed that Tfam engages the membrane most frequently at curved regions **(Fig. 2D)**, implying that the frequency of Tfam appearance at the membrane surface is determined by curvature.

The collective behavior of Tfam on membrane was further analyzed using reconstituted supported bilayer systems. Building on our previous work using tethering strategies to reconstitute dynamic IMM structures[33, 34], we developed double-stranded DNA-tethered lipid bilayers (DLBs)[35] with POPC and 20 mol% CL **(Fig. 2E)**. In this system, the intrinsic curvature of the membrane can be systematically adjusted by altering the length of the underlying DNA tethers, as confirmed by both Scanning Ion Conductance Microscopy (SICM) **(Fig. 2E)** and Atomic Force Microscopy (AFM) **(Extended Fig. 2A, B)**. The integrity and biophysical properties were validated by single-molecule tracking analysis **(Extended Fig.2C, Extended Movies 3 and 4)**. While probe molecules on the DTB were able to access the entire observation window, their motion deviated from the predominantly Brownian diffusion characteristic of typical solid-supported bilayers. Instead, we observed localized, obstructed diffusion, evident from both speed map and mean square displacement (MSD) analyses**(Extended Fig. 2E, F)**. This finding aligns with previous works on other tethering systems[36, 37], suggesting heterogeneities in lipid packing. Notably, these effects were significantly enhanced in the presence of cardiolipin, consistent with our molecular dynamics (MD) simulation results.

Tfam labeled with Alexa 488 was added to the bilayer surface, initiating protein enrichment at concentrations as low as 30 nM **(Fig. 2C)** without additional crowding reagents. Confocal Z-reconstruction**(Extended Movie 5)** revealed that these aggregates extend into the solution, indicating that protein clustering is not confined strictly to the 2D membrane surface. Fluorescence recovery after photobleaching (FRAP) showed partial recovery **(Fig. 2C)**, which remained consistent regardless of systematic Tfam concentration. Quantification of the diffusion coefficient of Tfam within condensates using fluorescence correlation spectroscopy (FCS) confirmed significantly slower apparent diffusion compared to Tfam in solution at the same concentration **(Extended Fig. 3C)**. Collectively, these data provide strong evidence that Tfam undergoes a phase transition at the surface of IMM mimetics, forming liquid surface condensates.

Since the recruitment of DNA by Tfam condensation is considered a key principle of nucleoid organization[10, 20], we tested if condensates formed at the membrane surface retained this capability. We introduced two different mtDNA fragments(ND1, and ATPase6) into the Tfam-IMM mimetic system. In the absence of Tfam, or at low concentrations where no condensates formed, the negatively charged DNA fragments were not recruited to the bilayer surface. However, upon the formation of Tfam surface condensates on the bilayer, we observed strong colocalization of both fractions with the Tfam condensates **(Fig. 2H,I)**. This indicates that Tfam surface condensates formed on the bilayer retain the ability to sequester mtDNA, similar to three-dimensional Tfam condensates formed in solution[10, 20]. Crucially, our data suggests that membrane-proximal Tfam condensates can effectively capture DNA, establishing a potential prerequisite step for nucleoid formation at the inner membrane.

### Tfam Prewets at the IMM Mimetic Surface and Show Multiphase Coexistence Feature

The low concentration of Tfam as we observe the condensation formed on bilayer surface follows one of the key criterial of surface prewetting (**Extended Figure 3A**). To identify the key physical factors controlling this membrane-associated Tfam assembly, we investigated the dependency of ionic strength on Tfam condensation with the existance of IMM mimetic surface under different Tfam concentrations. Under physiological ionic strength (150∼300 mM NaCl), surface condensates were observable with Tfam concentrations in the buffer as low as 30 nM. The assembly process showed a clear inverse correlation with ionic strength; elevating NaCl concentration to 500 mM–1M progressively suppressed condensate formation (**Fig. 2E**). This salt sensitivity implicates multivalent electrostatic interactions remains key driving forces governing this membrane-associated LLPS proccess, consistent with Tfam condensation in solution.

Atomic Force Microscopy (AFM) provided orthogonal structural validation of Tfam condensation on the bilayer surface. In the absence of protein, CL-DLBs displayed a relatively smooth surface topography characterized by intrinsic roughness containing both positive and negative curvature (**Fig. 3C**). Upon the addition of Tfam, we observed the emergence of discrete nanoscale patches with a height of approximately 10 nm (**Fig. 3I, 3M, Extended Fig. 3B**). The area of these membrane-associated condensates increased progressively as Tfam concentration was raised to 120 nM. Notably, the observed height (10 nm) represents a thin, surface associated molecular film, rather than the large spherical condensates typical of bulk phase separation. This restricted spreading at z-axles is a hallmark of the prewetting regime, confirming that the membrane surface induces a quasi-2D condensate distinct from 3D bulk separation.

**Fig. 3.**
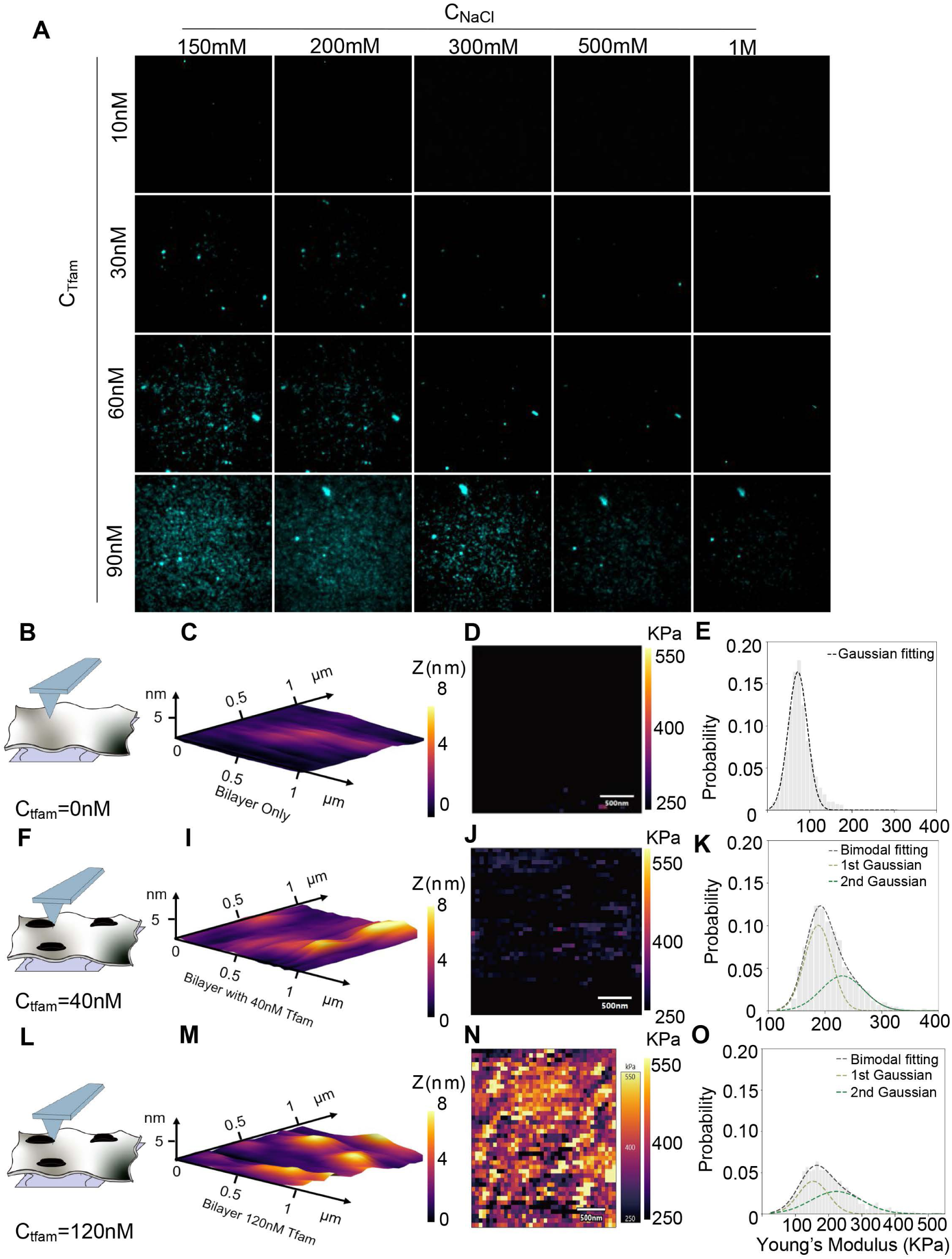
Characteristic of Tfam Surface condensates. **A.** Tfam condensate at the membrane surface is both concentration and electrostatic dependent. Condensates form above ∼30 nM at physiological salt but are progressively suppressed at ≥ 500 mM NaCl. Each panel represents a 25 × 25 *µm*^2^ region. **B.** AFM tapping of bare CL-DLBs and **(C)** with 40 nM **(F)**and 120 nM **(L)** Tfam concentration on bilayer. Z scale readout of the thickness suggesting Tfam condensate on the bilayer are proximately 2-3 layers of molecules**(I, M)**. whereas distribution of young’s modulus distribution of bared bilayer **(D)** and Tfam with either 40 nM **(J)** and 120 nM **(N)** bulk concentrations, suggest the mechanical properties of Tfam undergoes significant transition with elevated Tfam concentration which is summarized by bimodal fitting of correspondence young’s modulus mapping of bared bilayer **(E)**, with 40 nM Tfam**(K)** and with 120 nM Tfam **(F)**, correspondingly.

Mechanical mapping of the condensates revealed a striking concentration-dependent change in mechanical properties. The pristine lipid bilayer (0nM Tfam) exhibited a uniform mechanical response, characterized by a single Gaussian distribution of Young’s modulus with an average of approximately 72.00 kPa (**Fig. 3E)**.

In contrast, Tfam addition introduced a marked shift. Both 40 nM and 120 nM Tfam samples showed a broader Young’s Modulus distribution across the observed area, with distinct regions of higher stiffness evident in the corresponding mapping results (**Fig. 3J and 3N**). These mechanical distributions were further analyzed by bimodal Gaussian fitting (**Fig. 3K, 3O**). The bimodal fitting revealed that the Young’s Modulus distributions for both 40 nM and 120 nM Tfam concentrations consistently exhibited a high Young’s Modulus component that was approximately double the value of the lower modulus component. At the higher Tfam concentration (120 nM), the relative proportion of this high Young’s Modulus component was significantly elevated compared to the 40 nM concentration, and this component also showed a broader distribution (**Fig. 3O vs. 3K**).

Crucially, even the first, softer component of these bimodal fits did not perfectly align with the Young’s Modulus of the Tfam-free bilayer. This observation suggests that even regions not forming distinct condensates are mechanically perturbed, or that Tfam forms a distributed, less stiff phase that is difficult to completely separated from the bare membrane’s mechanical properties.

### A Thermodynamic Framework for Curvature-Mediated Prewetting

The emergence of surface-bound Tfam assemblies from an undersaturated bulk solution implies that the membrane is thermodynamically involved, functioning as more than merely a structural scaffold. To rationalize protein condensation behavior we observed, we introduce a minimal thermodynamic framework based on the Cahn theory of wetting [38]. We define the system using a scalar order parameter, *ϕ*(*x, y, z*), representing the local protein volume fraction. The Helmholtz free energy functional, F[*ϕ*], incorporates a bulk contribution describing fluid-fluid demixing and a surface free energy density *f_s_* at the membrane interface:

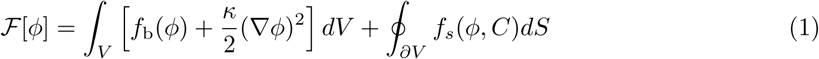

Here, *f*_b_ represents the standard interaction energy (e.g., Flory-Huggins), while the surface term *f_s_* encodes the protein-membrane interaction. Crucially, we employ a local thermodynamic approximation where the membrane curvature *C*(*x, y*) acts as an geometric cue. Based on our MD simulations (see **Fig. 2C and 2D** for details) showing curvature-driven lipid sorting, we propose that the local curvature acts as an *effective external field*, *h*(*C*), which modulates the surface chemical potential. Specifically, regions of negative curvature (*C <* 0) create a preferential binding potential, *µ*_eff_ ≈ *µ*_0_ +*ηC*, effectively lowering the local energy barrier for surface accumulation (see details in SI). Our model predicts that membrane curvature heterogeneity modulates the local TFAM–membrane binding affinity, giving rise to intrinsic heterogeneity in surface accumulation even in the absence of macroscopic condensation**(Fig. 4A)**. As a consequence, the membrane separates into two distinct surface states, referred to as thin and thick phases, corresponding to regions with different local curvature and binding environments**(Fig. 4G and 4I)**. This theoretical prediction finds compelling support in our atomic force microscopy (AFM) measurements **(Fig. 3E, 3K, 3O)**: the lower Young’s Modulus component obtained from bimodal fitting of Tfam-treated membranes is significantly higher than the elasticity of the pristine membrane, suggesting the formation of a distributed Tfam-associated layer (the ’thin phase’). Simultaneously, a distinct, stiffer component emerges, corresponding to the assembly of a thicker, more elastic condensed layer (the ’thick phase’).

**Fig. 4.**
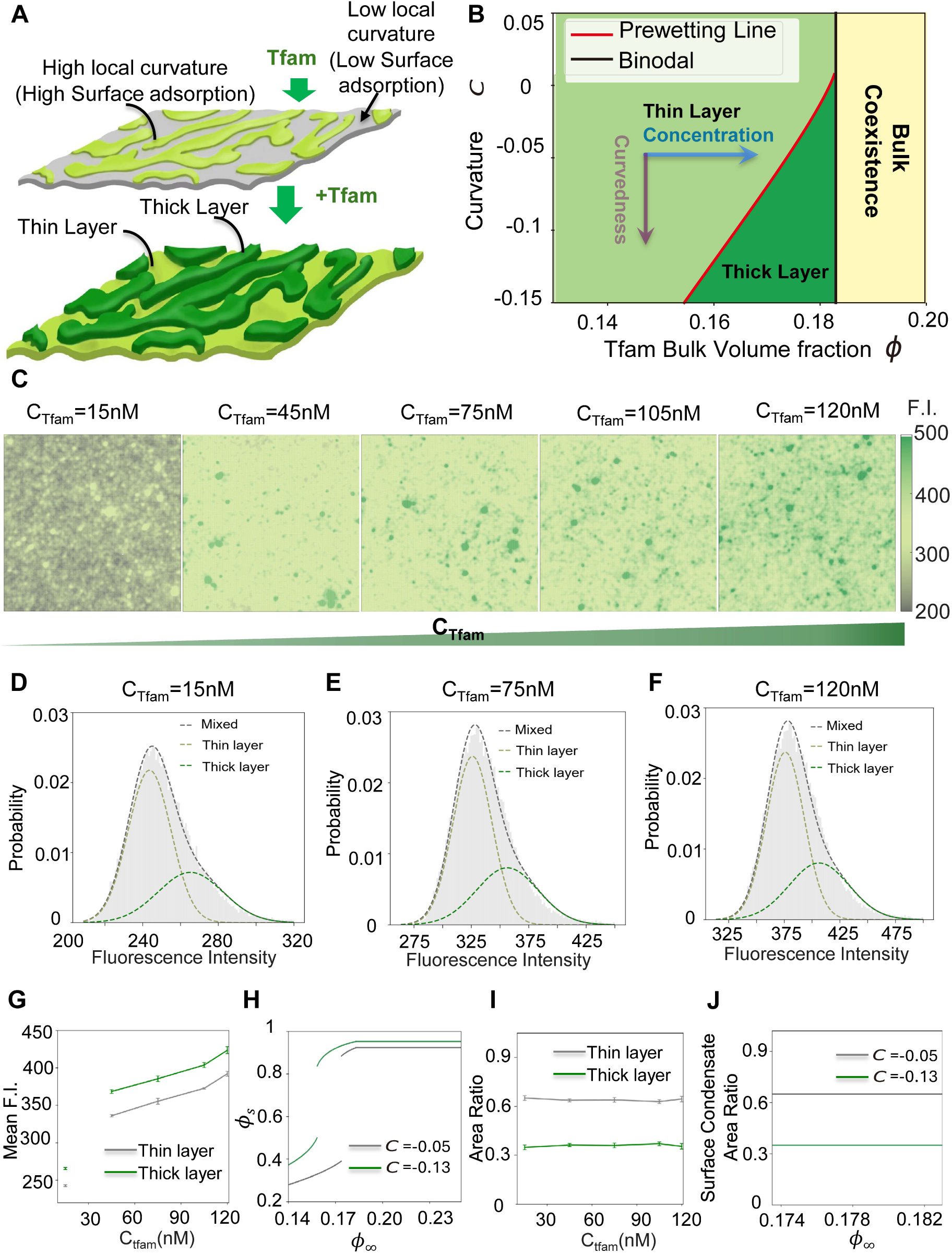
Thermodynamic model of curvature-driven prewetting. **(A)** Schematic of Tfam adsorption biased by membrane curvature. **(B)** predicts that a stable thick condensate forms above specific thresholds of negative curvature (purple arrow) and bulk concentration (blue arrow), transitioning from a thin adsorbed layer. (**C**) Intensity profile obtained from TIRF microscopy showing Tfam accumulates on the bilayer with phase coexistance (area shown 32×32 *µm*^2^). **(D-F)** Examplary intensity profiles of Tfam at the bulk concentration of 15 nM **(D)**, 75 nM **(E)** and 120 nM **(F)** with biomodal fitting indicating thin (light green Gaussian curve) and thick (dark green Gaussian curve), with the statistic results are summarized in **G** (mean fluorescence intensity) and **I** (area ratio of the thin and thick layers). The theoretical simulation results based on out thermodynamic model are summarized in **H**, and **J**. All error bars representing standard deviation from at least three biological replicates.

To further quantify this curvature induced surface hetergeneity,without measuring the signals from lipid bilayer, we experimentally characterized it using quantitative fluorescent measurements of Alexa 488 labeled Tfam titrated on IMM-mimetic membranes (**Extended Movie 6**). Statistical analysis of the fluorescence intensity distributions from TIRF microscopy using a bimodal fitting procedure revealed the coexistence of two surface phases across a broad range of bulk TFAM concentrations (**Fig. 4C-4F, Extended Fig. 3A**). This bimodal character persists at all concentrations, demonstrating that surface heterogeneity precedes(**Fig. 4H**) and shapes subsequent condensation behavior rather than emerging as a result of it.

Upon increasing the bulk TFAM concentration, both surface phases independently underwent surface condensation, giving rise to condensed assemblies with distinct characteristic intensities. This behavior reflects the difference in local binding affinity between the two phases and indicates that surface condensation proceeds along two parallel pathways set by membrane curvature. Quantitative bimodal analysis confirm this picture (**Fig. 4F**): while the relative populations of the two surface phases remain approximately constant at low concentrations, increasing TFAM concentration selectively promotes condensation within each phase. The properties of thin, thick layer were further quantified by fluorescence correlation spectroscopy **(Extended Fig. 3C, Extended Fig. 3D)**.

These observations are consistent with the theoretical prediction of a prewetting transition, in which surface accumulation undergoes a discontinuous transition from a microscopic adsorbed state to a macroscopic condensed film [11, 38, 39]. Importantly, in the present system this transition occurs on a heterogeneous membrane background, resulting in curvature-dependent condensed surface states. The experimentally measured normalized fluorescence intensities are in qualitative agreement with the thermodynamic framework detailed in the SI **(Fig. 4G-4J)**. Specifically, the experimental evolution of the mean fluorescence intensity for both phases **(Fig. 4G)**closely mirrors the theoretical prediction for the surface order parameter *ϕ_s_***(Fig. 4H)**. In both experiment and theory, the density of the identified thin and thick layers increases with rising bulk concentration, consistent with the progressive filling of available surface states.

Moreover, a key signature of this curvature-dictated mechanism is the spatial stability of the phases. Our experiments revealed that the relative area ratio between the two surface phases remains constant, independent of the bulk Tfam concentration (**Fig. 4I**). This robustness is perfectly reproduced by our model (**Fig. 4J)**, which predicts fixed surface coverage fractions. This confirms that the spatial extent of the condensed phases is predetermined by the intrinsic curvature landscape of the membrane template (the slots available for binding), rather than being driven by the lateral growth of domains as typically seen in flat membrane scenarios.

### Curvature-Mediated Energetic Barrier Lowering

Our thermodynamic model predicts that the transition from a thin adsorbed layer to a thick condensate depends not only on bulk concentration but critically on membrane curvature (**Fig. 4B**, **Fig. 5A**). To rigorously validate this prediction, we engineered three distinct bilayer architectures by manipulating both chemical composition and tethering geometry. **Type I** membranes (POPC tethered with 80BP DNA) serves as controls with minimum curvature. **Type II** bilayers (POPC with 20% CL with 80BP-DNA tether) in principle elevated membrane curvature due to CL’s unique feature on negative curvature, which was the dominant IMM mimetics applied in the previous experiments. Finally, to introduce significant macroscopic curvature, we developed **Type III** bilayers using the same IMM composition but tethered with a 1:1 mixture of 40 and 80 BP DNA strands.

**Fig. 5.**
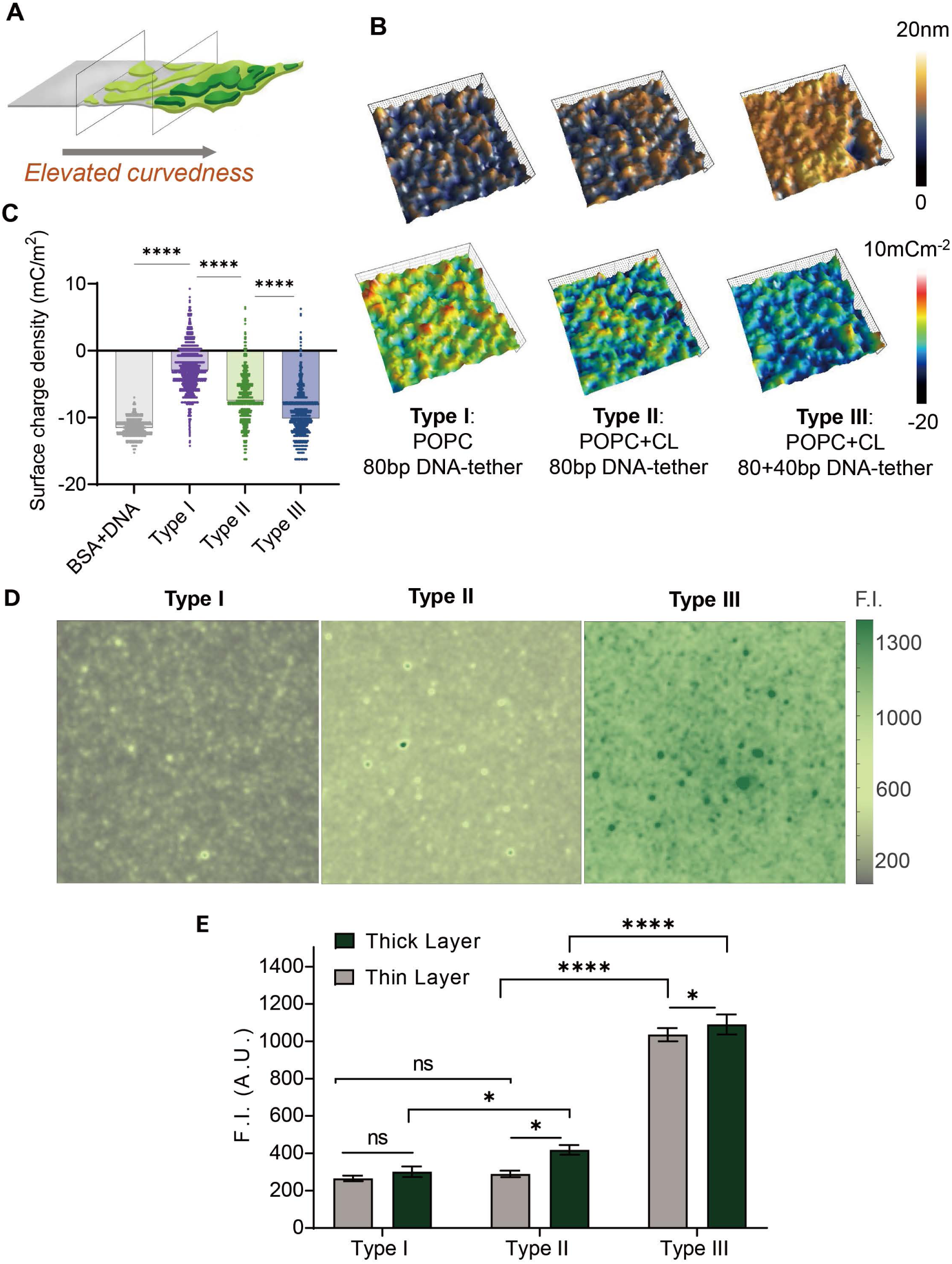
Membrane curvature promotes Tfam prewetting. **(A)** Schematic illustrating the curvature-dependent transition from a thin adsorbed layer to a thick condensate. **(B)** Topographical characterization of three engineered lipid bilayers (Types I-III) using SICM. Top row displays height maps; bottom row shows corresponding surface charge distribution. (**C**) Quantification of average surface charge density across the three bilayer types. **(D)** Represnetaitve TIRF images and segmented results (area shown 32×32 *µm*^2^). **(E)** Fluorescence intensity of Tfam on Type I, II and III bilayers analyzed from bimodal fitting. Statistic analysis were conducted using one-way ANOVA test from at least three independent replicates.

We simultaneously resolved the surface topography and local charge density of these systems using Scanning Ion Conductance Microscopy (SICM) **(Fig. 5B)**. While the underlying substrate displayed a homogeneous negative charge, the decoupling provided by the DNA tethers resulted in a overall neutral charge distribution for **Type I** bilayers. However, the introduction of geometric complexity in **Type II** and **Type III** systems fundamentally altered the surface landscape (**Fig. 5B upper row**). **Type III** bilayers, characterized by the highest surface roughness (**Extended Fig. 2A, 2B**), exhibited most significant local accumulations of negative charge.

This membrane heterogeneity was further elucidated by lipid diffusion analysis. Mean-squared dis-placement (MSD) and speed maps of lipid probes (TR-DHPE) revealed that regions of high curvature and charge coincided with obstructed lipid diffusion (**Extended Fig. 4A-C**). These findings establish a clear biophysical hierarchy: engineered membrane curvature with local ”high affinity patches”, echos our MD results and preset conditions in our thermodynamic model.

To define the relationship between membrane curvature and phase behavior, we performed comparative TIRF titration experiments (**Extended Fig. 5**). Visual inspection of Tfam on bilayer at comparable concentration (e.g., 120nM) reveals a dramatic hierachy (**Fig. 5D**): while **Type I** (low curvedness, neutral) and **Type II** (chemical enhanced intermediate curvature) surface display relatively low fluorescence, **Type III** bilayers (macroscopic curvature) exhibit bright dense distinct condensates.

Quantitative analysis of the fluorescence intensity (**Fig. 5E**) confirms a non-linear jump in the phase density, mirroring the prewetting transition (red line) along with the variation of membrane curvature predicted by our phase diagram (**Fig. 4B** ). In specific, on **Type I** control bilayers, the mean fluorescence intensity remains uniformly low (∼260 A.U.), with no statistic significance between the thin and potential accumulation sites. This confirms that on membrane with relatively low curvature, the local chemical potential (*µ*_eff_) is insufficient to overcome the nucleation barrier, effectively trapping the system in the dilute prewetted states. **Type II** bilayers, as illustrated in **Fig. 4C** with the highest concentration distribution summarized in **Fig. 5E**, present an intermediate thermodynamic state: chemical affinity initiates weak condensation, however, without the macroscopic topological support, phase remains tenuous and was not able to reach high density.

The critical transition is triggered by macroscopic curvature in **Type III** membranes, here the synergistic combination of chemical and geometric roughness drives a massive amplification in protein density during nucleation. The fluorescence intensity of the condensate phase surges to higher than 1000 A.U., nearly three-fold increase compare to **Type II** and four fold compared to **Type I** (**Fig. 5E**). Such ”intensity jump” we observed during our experimental model, corresponds directly to the vertical shift along the Y-axis of the theoretically predicted phase diagram (**Fig. 4B**). Comparing to **Type I** and **II** membranes, the high curvature of **Type III** membranes lowers the effective chemical potential sufficiently to bridge the energy gap. This allows the system to ”jump” into the stable, high-density ”liquid” regime. Thus, membrane curvature does not merely localize the protein; it fundamentally alters the thermodynamic state, enabling the transition from a weak surface coat to a functional, high-density nucleoid.

### Membrane Curvature Dictates Nucleoids Assembly In Situ

Our thermodynamic model predicts that Tfam phase separation is universally regulated and spatially templated by membrane topography. This implies that Tfam condensates should respond predictably to any cue that alters IMM geometry, regardless of the upstream trigger. To test the dominance of this geometric regulation, we challenged the system with MPP+, a complex I inhibitor that induces mitochondrial swelling and cristae unfolding (**Fig. 6A, Extended Movie7**). Low-dose MPP+ treatment allowed us to chemically flatten the membrane without genetic manipulation, and then test whether curvature rescue via Opa1 overexpression, either prophylactically or therapeutically, could restore condensation.

**Fig. 6.**
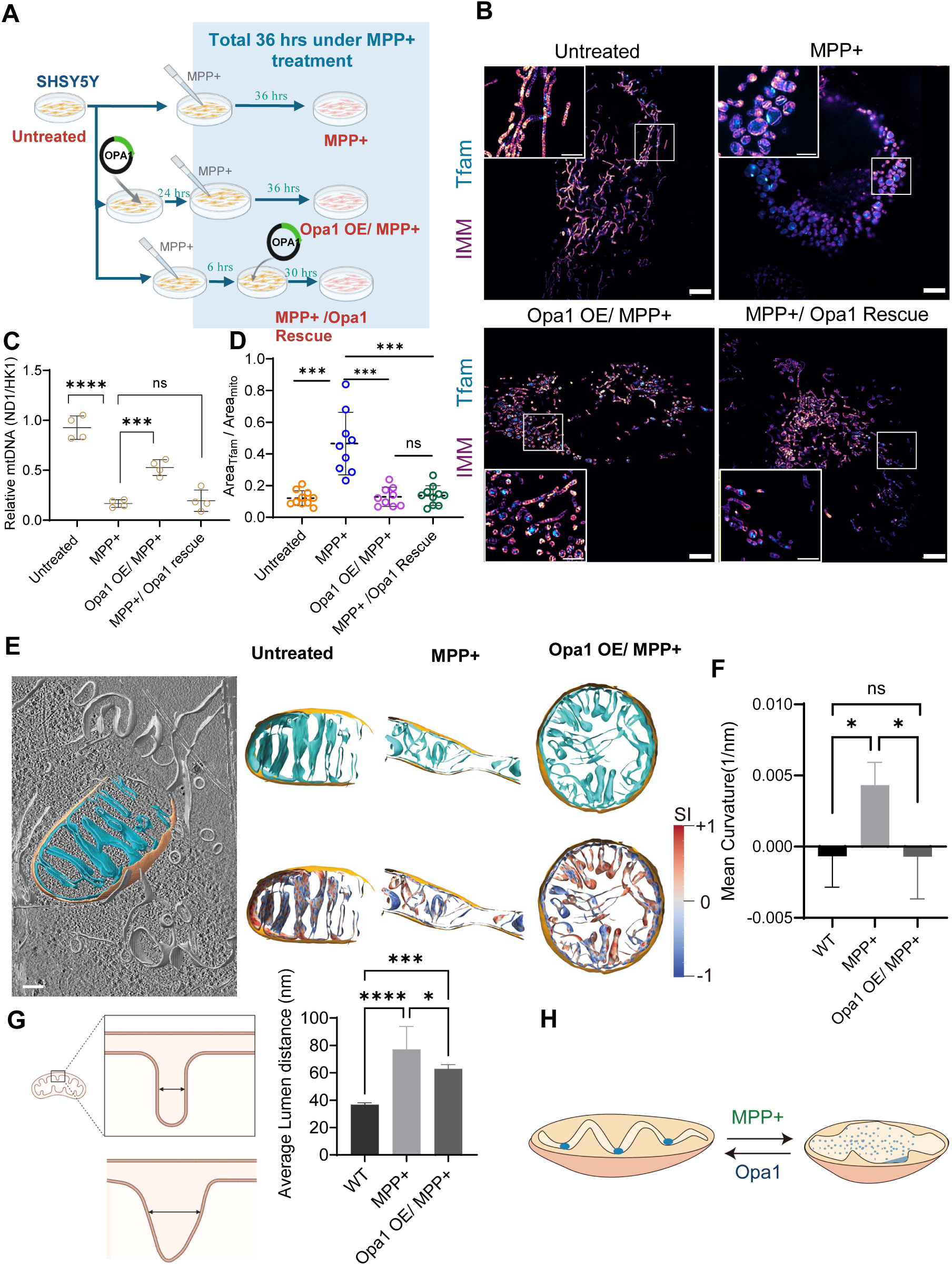
In situ correlation between IMM curvature and Tfam organization. **(A)** Experimental schema for chemical disruption of IMM structure using MPP^+^ in SH-SY5Y cells, with prophylactic or curative Opa1 overexpression (OE).**B** SIM imaging comparing Tfam distribution. MPP^+^ treatment (36 h) induces mitochondrial swelling and Tfam dispersal, while Opa1 OE restores discrete nucleoid structures. (Scale bars: overview 5 *µ*m, enlarged, 2 *µ*m). (**C**) Analysis of mtDNA levels versus Tfam packing fraction. Notably, Opa1 OE restores Tfam condensation despite persistently low mtDNA levels, effectively decoupling nucleoid structure from DNA content. (**D**) Quantification of Tfam matrix occupancy (Area_Tfam_/Area_mito_).**(E)** Representative cryo-electron tomogram of a single mitochondrion (left; cyan: inner membrane, orange: outer membrane; scale bar: 100 nm). Right panels show 3D segmentation (top) and IMM Shape Index (**SI**) mapping (bottom).**(F–H),** Quantification showing that Opa1 OE recovers IMM negative curvature (**F**) even while cristae distance remains enlarged (**G**). This curvature recovery directly correlates with restored Tfam packing (**H**). Statistics: One-way ANOVA, n≥ 3; ∗*P<*0.05, ∗ ∗ *p<*0.01, ∗ ∗ ∗*P<*0.001, ∗ ∗ ∗ ∗ *P<*0.0001*, ns* : not significant).

We correlated the nanoscale geometry of these perturbation with Tfam Phase behavior using a combination of Cryo-electron tomography (CryoET) **Extended Movies 8-16** and live cell SIM. CryoET results of MPP+ treated mitochondria revealed a profound alternation in membrane curvature. While untreated cells were dominated by lamellar cristae enriched in negatively curved and saddle-shaped regions(Shape index *<* 0), MPP+ treatment shifted the IMM toward an average positive curvature (convex) as a result of matrix swelling and vesiculation (**Fig. 6E, 6F**). Quantitatively, positively curved regions expanded to occupy 40.1% of IMM surface, compared to only 29.1% in controls (**SI**).

Crucially, this geometric inversion directly dictated Tfam phase dynamics. Live-cell SIM imaging revealed that Tfam in these swollen, positively-curved mitochondria lost its punctuated structure and dispersed throughout the matrix, with occasional enrichment close to the membrane surface (**Fig. 6B**, upper row). This observation was confirmed by live cell- STED imaging where Tfam average intensity was confirmed lower than those in untreated cells. In addition, fluorescent lifetime analysis under STED illumination showed that the fluorescence lifetime of Tfam increased to match that of monomeric mNeogreen, indicating a loss of molecular packing and self-quenching[40](**SI Fig. 2E**). Thus the loss of negative curvature shifts the ”prewetting” boundary condition, prevents thick layer formation, forcing more Tfam into the bulk, diluted phase.

The Opa1 rescuing assays were applied to test the causality. Live cell SIM showed under both conditions, Tfam distribution were fully recovered (**Fig. 6B**, lower row) regardless of the ongoing chemical stress. Interestingly, under Opa1 rescue conditions, mtDNA level were similar to MPP+ treatment, regardless of the clearly recovered nucleoid structure, suggesting Tfam condensation is not dependent on mtDNA level. Consider both conditions showed similar effects on Tfam distribution, CryoET analysis were conducted on Opa1 OE/MPP+ cells (**Fig. 6E**). Quantification under this condition revealed the structural basis for recovery: while Opa1 did not fully reconstitute native cristae width (**Fig. 6G**), it successfully restored the balance of curvature states (**Fig. 6F**). The fraction of positively curved membrane was reduced to 30.5%, and the landscape was again dominanted by saddle and negative geometry (**SI**).

The tight correlation between these *in situ* structural and live cell imaging establishes a clear biophysical cue: membrane curvature is the proximal regulator of nucleoid organization. The transition from condensed to dispersed Tfam tracks specifically with the shift from negative/saddle to positive curvature(**Extended Data 7**), independent of cristae lumen width or mtDNA copy number. This confirms that the membrane landscape acts as a thermodynamic mold, where local curvature defines the spatial boundaries for Tfam condensation.

## Discussion and conclusion

Physical principles governing cellular organization extend across scales, from the nanoscale assembly of protein complexes to the microscale compartmentalization of organelles[4, 41, 42]. While alternmations in membrane geometry are fundamental to membrane fusion and fission[43, 44], the role of membrane morphological dynamics in regulating biomolecular organizations , specifically through the potential biomolecular condensations, has remained elusive. Here, we demonstrate that IMM curvature facilitates the organization of the mitochondrial nucleoids not simply by squeezing bulk 3D condensates into a small volume, but through a previously unreported thermodynamic mechanism: geometry-directed prewetting. Importantly, simple geometric confinement cannot overcome the massive thermodynamic energy barrier required for Tfam condensation at low protein concentration limits we observed, without the addition of artificial crowding agents. Instead, in the context of the Cahn wetting framework, the mesoscale topography (macroscopic curvature) acts as a localized thermodynamic field that explicitly modifies the surface free energy landscape.

By coupling local curvature to the effective chemical potential *µ*_eff_, the negatively curved membrane dramatically lowers the nucleation barrier (Δ*G*^∗^). It shifts the prewetting coexistence boundary, actively driving a non-linear transition from a dilute thin adsorbed layer to a locally enriched ”thick” condensation layer with the condition where the bulk concentration is orders of magnitude below the bulk saturation concentration (*C*_sat_). Thus, the membrane geometry does not merely corral the biomolecular condensates such as nucleoids, but acts as a thermodynamic switch that enables and spatially templates phase separation under strick physiological boundaries.

Our model fundamentally distinguishes Tfam from canonical curvature-sensing proteins[45]. Classical sensors, such as BAR domains, rely on shape complementarity to recognize bent membranes[46, 47]. In contrast, our data identify Tfam as a thermodynamic sensor: it detects the modified free energy landscape created by membrane curvature. In the IMM, curvature drives lateral lipid heterogeneity, specifically enriched cardiolipin at negative and saddle geometries due to its conical shape. The sorting creates local high affinity patches that lower chemical potential for the thick layer of Tfam to condensate. Consequently, Tfam undergoes a prewetting transition and form a dense, stable phase that is restricted by membrane curvature. This mechanism allows the cell to utilize the membrane not just as boundary, but participate in regulating spatial organization of biochemical activity.

Crucially, our in situ perturbation experiment identify membrane curvature is one of the dominant parameter for this transition, acting universally regardless of the upstream biological trigger. The chemical reagent induced flattening of the membrane induces ”dewetting” transition, which could be reversely transition by overexpressing structure proteins which restores the curvedness and overall negative curvature of the IMM. This tight correlation establishes that the specific balance of membrane curvature states defines the thermodynamic boundary conditions for nucleoid integrity.

This regulatory axis offers a physical lens for understanding mitochondrial diseases. For example, in Barth syndrome, defective cardiolipin remodeling may disrupt nucleoid organization[48] not by merely altering lipid chemistry, but by failing to generate the curvature-dependent energy gradients required for the prewetting transition. Similarly, the loss of cristae-shaping factors (e.g., Opa1) during aging[49, 50], or membrane swelling induced by toxins[51], leads to genome instability precisely because the geometric boundary conditions for condensation are removed. The dispersion of Tfam we observed upon curvature loss suggests that maintaining cristae morphology is not just structural, but a thermodynamic necessity for localized spatial organization of the genome.

The complexity of membrane curvature in cellular membrane were evolving with cellular function. However, the mechanism echos evolutionary heritages. Bacteria, the ancestors of mitochondria, also position their nucleoids in close association with the inner membrane, which enables the partitioning of genetic material during cell division[52, 53]. The reliance on geometry to sculpt lipid heterogeneity and modulate phase behavior suggests an evolutionarily conserved strategy. Potentially by exploiting the physics of wetting, cells achieve exquisite spatial control over membrane enclosed materials without the energetic cost of active transport motors.

Overall, our study establishes geometry-directed prewetting as a fundamental mechanism by which cellular architecture controls biomolecular spatial organization. By demonstrating that membrane curvature acts as a thermodynamic control parameter for protein condensation, we reveal that organelle membranes function as active regulators rather than passive boundaries. The mitochondrial inner membrane exemplifies this principle: its complex topography creates spatially heterogeneous energy landscapes that template phase separation at specific geometric regions, achieving precise subcellular organization through the physics of wetting. This curvature-mediated mechanism transcends the specific biology of mitochondria, suggesting that surface-catalyzed condensation represents a universal organizing principle governing phase behavior in membrane-rich environments from organelles to synthetic compartmentalized systems.

## Methods

### Cell culture and treatments

SH-SY5Y cells were maintained under undifferentiated conditions in Dulbecco’s modified Eagle’s medium (DMEM; Gibco) supplemented with 10% heat-inactivated fetal bovine serum, 2 mM glutamine and 1% penicillin–streptomycin. Mouse thymic epithelial cells (mTECs) were cultured in DMEM supplemented with 10% fetal bovine serum and 1% penicillin–streptomycin. HEK293T cells used for lentiviral production were cultured under standard DMEM conditions. All cell lines were routinely tested and confirmed to be free of mycoplasma contamination. For MPP^+^ treatment, SH-SY5Y cells were exposed to 500 *µ*M1-methyl-4-phenylpyridinium (MPP^+^; Innochem) for the indicated durations. Full-length human OPA1 isoform 1 was expressed from a pcDNA3.1 backbone at time points specified in Fig. 6.

### Plasmid construction

Full-length human Tfam (UniProt ID Q00059) DNA sequence was synthesized by GENEWIZ. For live-cell imaging, Tfam (amino acids 1–246) was fused at its C terminus to mNeonGreen via a flexible glycine–serine linker (GGGGSGGGGS) and cloned into pcDNA3.1(+). For protein expression, Tfam without the mitochondrial targeting sequence (amino acids 43–246) was codon optimized for E.coli expression by Xianhua Biotech, tagged with C-terminal hexahistidine and cloned into a pET-28a vector.

### Stable cell line generation

Stable SH-SY5Y cell lines expressing Tfam–mNeonGreen were generated by transfection using Lipofectamine 2000 (Invitrogen) followed by selection with G418 (Beyotime). Expression was verified by live-cell fluorescence imaging. Stable cell lines were used within five passages after selection.

Opa1 knockout mTECs were generated using a lentiCRISPRv2 system. Paired single-guide RNAs targeting Opa1 (Supplementary Table 2) were cloned into the lentiviral backbone and packaged in HEK293T cells using psPAX2 (Addgene #12260) and pMD2.G (Addgene #12259). Viral supernatants were collected 48 h post-transfection and used to transduce target cells in the presence of polybrene (YEASEN). Transduced cells were selected with puromycin, and single-cell clones were validated by genomic sequencing and immunostaining (Fig. 1).

### Tfam protein expression and purification

Tfam was expressed and purified by Xianhua Biotech based on previously described protocol [54] with minor modifications. Briefly, cells were cultured in LB media at 37 ^◦^C, and induced by 0.1 mM IPTG at OD 0.5 overnight. Purification included Ni-affinity chromatography in Tris buffer containing 500 mM NaCl, followed by size-exclusion chromatography on a Superdex 200 Increase column (Cytiva) in buffer containing 25 mM Tris and 300 mM NaCl (pH 7.4). Tfam fractions were collected, concentrated and subjected to heparin affinity chromatography to remove bound nucleic acids. Purified protein (2 mg*/*mL) was labeled with Alexa Fluor 488 NHS ester (Invitrogen) at a dye-to-protein molar ratio of 1–2.

### Live-cell super-resolution imaging and analysis

For live cell mitochondrial imaging, the IMM was labeled with PKmito Orange according to published protocols. Of which, Structured illuminated microscopy (SIM) images were aquired using a His-SIM (CSR Biotech) build on an IX83 inverted microscope (Evident), equipped with live cell incubation chambers (Tokaihit). In specific, an Olympus objective (100×, NA 1.45 oil objective) was applied for data acquisition. All SIM images were acquired under preset model 2D-SIM2 using both 488 nm and data were reconstructed using Finer (CSR Biotech), where both Wiener reconstruction and background substruction were conducted according to previously published method.

Live cell STED microscopy was conducted using Abberior Facility Line microscope (Abberior Instruments) fluorescence microscope built on a motorized inverted microscope IX83 (Evident) equipped with Olympus 60× objective (NA1.42, oil immersion) and live cell incubation chamber.

In specific, Tfam-mNeogreen signals were imaged using a pulsed 485 laser (1 mW pulsed 40 MHz), 595 nm high power laser module was applied as STED laser module with both automatic beam size control equipped in the system. For PKmito Orange labeling of IMM was imaged using a combination of fluorescence excitation module with 561 nm laser combined with 775 nm STED laser. The fluorescence lifetime analysis under STED illumination was enabled by the TIMEBOW software package implements in the Lightbox acquisition and analysis software, where the time-binned data from FLIM-capable array detection (MATRIX) and analyzed using GPU-based phasor analysis embedded in the module.

### Cryo-ET sample preparation, Cryo-FIB milling and data collection

Quantifoil R2/1 200-mesh Au grids were glow-discharged for 45 s s on both sides using a plasma cleaner, then coated with poly-L-lysine (PLL). SH-SY5Y cells were seeded onto the treated grids and cultured overnight in DMEM medium. Following experimental treatment, the grids were plunge-frozen in liquid ethane using an FEI Vitrobot Mark IV (Thermo Fisher Scientific) at 16 ^◦^C and 95 % humidity. The plunge-freezing process involved 2 s wait time and 9 s of one-sided blotting from the back of the grid using Whatman filter paper (Sigma-Aldrich) with a blotting force of 8. All subsequent handling and transfers were performed under liquid nitrogen.

The plunge-frozen grids were loaded into an Aquilos 2+ cryo-focused ion beam/scanning electron microscope (Cryo-FIB/SEM, Thermo Fisher Scientific). In the cryo-FIB chamber, the specimen was first sputter-coated with inorganic platinum for 30 seconds, followed by the deposition of an organometallic platinum layer for 40 seconds using an integrated gas injection system (GIS). This was followed by a 30-second inorganic platinum sputtering. During the FIB milling process, the specimens were kept at a near liquid nitrogen temperature to preserve their cryogenic state. The milling process was performed with a 30 kV Ga^+^ ion beam at a milling angle of 10°. The milling was carried out in several steps, progressively reducing the lamellae thickness using decreasing ion beam currents, ranging from 1 nA to 30 pA. The final lamellae thickness was reduced to approximately 160 nm. After milling, an additional platinum sputter coating was applied to improve the conductivity of the lamellae.

The Cryo-FIB milled grids were transferred to a Titan Krios cryo-transmission electron microscope (Thermo Fisher Scientific) for imaging. The microscope was equipped with a Selectris energy filter (slit width: 20 eV) and a Gatan BioContinuum K3 direct electron detector. Low-magnification search maps of lamellae were acquired at 11, 500× magnification, with clear mitochondrial features used as reference regions for data acquisition. Multiple tilt series were acquired at 33, 000× magnification (pixel size: 2.66Å) over a tilt range of −52° to 52° in 2° increments using a dose-symmetric acquisition scheme and super-resolution mode, as implemented in the Tomography 5.12 software (Thermo Fisher Scientific). The total electron dose during data collection was maintained between 120 − 140 e^−^*/Å*^2^, with the defocus set to −7*µ*m to optimize contrast.

### Tomogram processing, segmentation and quantification

Tilt series were motion-corrected and CTF-estimated in Warp 1.1.0. After removal of low-quality tilts, series were aligned using patch tracking and reconstructed by weighted back projection using AreTomo2. CTF deconvolution and missing-wedge correction were performed using IsoNet v0.2. Membranes were segmented using Membrain-seg and manually curated in Amira to annotate outer mitochondrial membrane (OMM), inner mitochondrial membrane (IMM) and other membranes. Surface meshes were generated and analysed using the Surface Morphometrics pipeline and PyCurv to extract curvature metrics (including curvedness, mean curvature and shape index) from vector- voting–based curvature analysis outputs. Tomograms and meshes were visualized in ParaView 5.13.1 and ChimeraX 1.9.

### Coarse-grained molecular dynamics simulations of Tfam with a curved membrane

The crystal structure of human mitochondrial transcription factor A (PDB 3TQ6) was used to model Tfam [32]. The coarse-grained (CG) model of Tfam was constructed using the Martini 2.2 force field [55]. The CG model of Tfam was prepared using the martinize.py program and subsequently embedded in an IMM-like membrane model consisting of POPC with 20 % cardiolipin (CL) via the INSANE protocol [56]. The ElNeDyn elastic-network approach was employed to restrain the protein structure, with a force constant of 1000 kJ*/*mol*/*nm^2^ and lower and upper limits of the cutoff distance set at 0.5 and 0.9 nm, respectively. The systems were solvated using a CG Martini water model and neutralized by adding NaCl at a concentration of 0.15 M to mimic physiological conditions. To prevent unwanted freezing, 10 % of the MARTINI water particles were replaced by the antifreeze water particles. To compensate for the known over-sticky issue in Martini 2.2, the force field parameters were tuned by increasing protein-water interactions by 10 %, as described in our previous work [57]. Electrostatic interactions were treated using a reaction field with a dielectric constant of 15. Non-bonded interactions were cut off at 1.1 nm. The simulations were performed with a 20 fs integration time step. The Verlet neighbor search algorithm was used to enable GPU-accelerated simulations. To investigate membrane curvature, a buckled lipid bilayer was constructed using the lateral compression method [31], resulting in a compression factor of 0.3. To maintain the curvature, the simulations were conducted in the NVT ensemble, with the v-rescaling thermostat used to maintain a temperature of 310 K. All atomistic and CG MD simulations were performed using Gromacs 2021.57. Four CG MD simulations were performed, each lasting 20 *µ*s.

### Fabrication of IMM mimicking DTBs

DTBs were prepared by vesicle fusion using small unilamellar vesicles (SUVs) on DNA-functionalized glass substrates. Glass coverslips (No. 1.5H, Marienfeld) were cleaned according to previously described protocol8, stored in 50% H_2_SO_4_ for at least 2 days and exposed to UV illumination for 10 min before use, with extensive ultrapure water rinsing between steps. Substrates were coated with 0.3 mg*/*mL BSA in TBS (2 − 6 h), incubated with 1.4 *µ*g*/*mL NeutrAvidin in TBS 20 mins, and then incubated with hybridized CHOL–DNA–biotin dsDNA (20 mins), followed by TBS rinsing. Bilayers were formed by fusion of SUVs. For single-molecule imaging, lipid mixtures contained 0.00001 mol% Texas Red–DHPE. SUVs were incubated on DNA-functionalized substrates for 30 min (room temperature or 37°C for cardiolipin-containing bilayers), followed by gentle rinsing to remove excess vesicles. For AFM validation, UV shadow-mask photopatterning was used to generate adjacent DNA-tethered and directly glass supported membrane regions on the same substrate for DNA supported height comparison.

### Optical settings for total internal reflection microscopy (TIRF) and fluorescent correlation spectroscopy (FCS)

Home assembled TIRF microscopy equipped with 488 nm, 561 nm and 647 nm laser (Coherent OBIS) with confocal based FCS module were applied for the single molecular characterization of lipid and protein diffusion in the reconstituted systems. In specific, TIRF module (FluoCa) and FCS module (Lightedgetech) were both mounted on IX73 microscope (Evident). TIRF images were collected using an Olympus objective (100×, NA 1.45 oil objective, Tokyo, Japan), and images were acquired using an EMCCD (Oxford instrument). While FCS data were collected using an 63× (NA 1.2, water immersion) Olympus objective.

### Tfam titration and Imaging

Low-power excitation settings were employed to minimize spectral bleed-through and oversaturation at high concentrations. Tfam was introduced into a 500 *µ*L imaging system at the designated concentration, with all additions made in 50 *µ*L aliquots and diluted with chamber solutions. Images were captured using a 488 nm laser set to 0.2 − 0.4 mW, with an exposure time of 10 − 20 ms and a gain of 500.

### Single molecule tracking, diffusion and membrane integrity analysis

Time-lapse total internal reflection fluorescence (TIRF) microscopy was used to track lipid and protein diffusion. Single-particle trajectories were analysed to extract step-size distributions and mean-squared displacement (MSD) profiles and were classified using anomalous diffusion models (Supplementary Methods). Trajectory fitting and analysis were performed using custom Python scripts adapted from the Octane single-particle tracking framework with user-defined parameter settings. Membrane integrity and mean square displacement was verified by summing over 1,000 frames of lipid tracks.

### Atomic force microscopy (AFM) characterization

Liquid-phase AFM imaging was performed to characterize membrane topography and mechanical properties before and after Tfam titration. Measurements were carried out on an AFM system (MFP-3D, Asylum Research) operating in AC mode for topography and contact mode for stiffness measurements with soft silicon nitride probes (SiNi, Budget Sensors, 0.27 N*/*m). Topographic maps and force spectroscopy data were acquired on DNA-tethered bilayers and supported membrane regions under physiological buffer conditions. Young’s modulus values were extracted from force–distance curves using a Hertz-model fit (Supplementary Methods).

### Confocal imaging and FRAP analysis

Confocal image of Tfam droplet on membrane surface and 3D scan was conducted using FV4000 confocal microscope (Evident), Images and analysis were conducted using 100×, NA 1.45 oil objective (Olympus). For Fluorescence Recovery after Photon Bleaching were conducted under the preset tornado bleaching model with fast bleach at the 30% laser power, and recovery was recorded accordingly.

### SICM imaging and surface charge analysis

All SICM images were recorded using the Dualscope 200 (Eaglenos Sciences, Inc.). Surface-charge quantification was performed according to the method described by Klausen et al [58]. Briefly, Quartz nanopipettes (inner radius ≈ 30 nm) were fabricated on a CO_2_-laser puller (P-2000, Sutter Instruments). Both the nanopipettes and the scanning bath contained the electrolyte solution (150 mM NaCl, 10 mM HEPES, pH 7.0). Supported lipid bilayers were scanned in constant-current mode at 25°C using a 1% set-point. Topographic images were acquired consecutively at +100 mV and −100 mV pipette bias. The pixel-wise height-difference map (Δ*d*) between the two bias images was converted to surface-charge density (*σ*). Images were plane flattened in SPIP (Image Metrology).

## Supplementary information

The details of thermodynamic modeling, supplemental methodology, all extended data and movies are included in SI.

## Acknowledgements

This work was financially supported by the National Natural Science Foundation of China Grants: 32350018 to Y.G. , 12501556 to X.Z., 22425704 and 82188101 to C.L., and 22225403 to D.L. . National Key Research and Development Program of China 2023YFF0722600 to Y.G., 2025YFA1308800 to C.L.; Zhejiang Provincial Natural Science Foundation of China LQN26A010001 to X.Z.; Shanghai Basic Research Pioneer Project (to Z.C), and Shanghai Key Laboratory of Aging Studies (19DZ2260400, to Z.C.); Shanghai Municipal Science and Technology Major Project to C.L., the Strategic Priority Research Program of the Chinese Academy of Sciences (Grant No. XDB1060000 to C.L.). Dr. Cong Liu is a SANS Exploration Scholar.

The authors thank Prof.James J. Chou from Shanghai Institute of Organic Chemistry, Prof. Luke H. Chao from Massachusetts General Hospital, and Prof. Paula P. Navarro from University of Lausanne for helpful discussions. We thank Dr. Tianyi Li from Abberior Co.Ltd, and Dr. Ji Tang for technical support on live cell STED and STED-FLIM. Y.W. thanks the Information Technology Center and State Key Lab of CAD&CG, Zhejiang University, for computational support.

## Declarations

X.H., L.S., and G.Z. contributed equally to this work. The conceptual framework was established by X.H., Y.G., X.Z., C.L., and Z.C. Experimental work and data analysis were conducted by X.H., L.S., and G.Z. Y.S., X.H., L.S., T.L., and J.C. prepared the cell lines and performed the biochemical analyses. Y.G., L.S., G.Z., and S.F. were responsible for the super-resolution microscopy and cryo-electron tomography (cryoET) data. Y.W. developed and simulated the molecular dynamics (MD) model. In vitro reconstitution experiments were carried out by X.H. and L.S. Y.Y. and S.G. performed atomic force microscopy (AFM) analysis, while Y.X. and D.J. conducted scanning ion conductance microscopy (SICM) analysis. Thermodynamic theories were formulated and the fluorescence titration signals were analyzed by Y.J. and X.Z. Instrumental resources were acquired, and the project was supervised by D.L., C.L., D.J., Y.G., and Z.C. The manuscript was written by Y.G., X.Z., X.H, L.S and G.Z.

Authors declare no conflict of interest.

## Notes

### Competing Interest Statement

The authors have declared no competing interest.

