## Supplementary material for "Curvature-mediated prewetting organize mitochondrial nucleoid": supplmental files

Supplemental Information

### Thermodynamic theory of prewetting on curved heterogeneous membranes

To elucidate the physical mechanism underlying the formation of TFAM surface condensates, we formulate a mean-field thermodynamic model. Distinct from classical wetting/prewetting on flat substrates, our experimental system exhibits spatial heterogeneity in membrane geometry (local curvature imposed by cardiolipin molecular shape and DNA scaffolds). To capture the interplay between these factors, we extend the classical wetting theory [1] to include a coupling between surface protein accumulation and local membrane curvature.

#### A. free energy functional with geometric coupling

We consider a semi-infinite system where the bulk solution (volume $V$) interacts with a membrane surface (area $\partial V$) at $z = 0$. The thermodynamic state is described by the bulk volume fraction profile of TFAM, $\phi(x, y, z)$. The membrane surface is treated as a quenched manifold characterized by one local field: the local mean curvature $C(x, y)$.

The Helmholtz free energy functional $\mathcal{F}[\phi]$ is given by:

$$\mathcal{F}[\phi] = \int_V \left[ f_b(\phi) + \frac{\kappa}{2}(\nabla\phi)^2 \right] dV + \int_{\partial V} f_s(\phi_s; C)\, dS. \tag{S1}$$

Here, $\phi_s \equiv \phi|_{z=0}$ denotes the surface concentration. $\kappa$ represents the interfacial free energy coefficient. The bulk free energy density $f_b(\phi)$ follows the Flory-Huggins solution theory:

$$f_b(\phi) = \frac{k_B T}{\nu} \left[ \frac{1}{n_b}\phi \ln\phi + (1-\phi)\ln(1-\phi) + \chi\phi(1-\phi) \right] + \omega\phi, \tag{S2}$$

where $\nu$ denotes the molecular volume of the solvent, $n_b$ represents the ratio of the solute molecular volume to that of the solvent, and $\chi$ characterizes the solvent–solute interaction parameter. The parameter $\omega$ denotes the coefficient associated with the internal free energy.

#### B. Curvature-Renormalized Surface Potential

The core of our model lies in the surface free energy density $f_s$, which must account for the fact that TFAM acts as a curvature sensor. We propose a surface potential that couples the protein concentration to the membrane curvature $C$:

$$f_s(\phi_s; C) = \mu_{\text{eff}}(C)\phi_s + \frac{1}{2} g \phi_s^2. \tag{S3}$$

Here, $g$ represents surface-enhanced cohesive interactions. The term $\mu_{\text{eff}}$ acts as a effective surface chemical potential, explicitly describing the driving force for adsorption:

$$\mu_{\text{eff}}(C) = \underbrace{\mu_0}_{\text{Basal}} + \underbrace{\eta C}_{\text{Geometric Affinity}}. \tag{S4}$$

The parameter $\mu_0$ denotes the chemical potential difference on a flat membrane. The geometric sensitivity is captured by the coupling $\eta C$. Here, $\eta$ is the curvature-coupling modulus, which quantifies how much the chemical potential is affected by the curvature. For $\eta > 0$, this term creates an effective potential well in regions of negative curvature ($C < 0$), driving local surface accumulation. This curvature-driven transport matches the physiological localization of TFAM condensates to the highly curved cristae of the inner mitochondrial membrane (see Fig. 5 in main text for details).

##### C. Equilibrium conditions

At equilibrium, the order-parameter profile $\phi(z)$ minimizes the Gibbs surface potential functional

$$\Omega[\phi] = \mathcal{F}[\phi] - \frac{\mu}{\nu_b} \int (\phi - \phi_\infty)\, dV,$$

where $\mu$ denotes the bulk chemical potential imposed by the reservoir.

Taking the first variation of $\Omega$ yields the bulk Euler–Lagrange equation together with the natural boundary condition at the membrane surface ($z = 0$). The latter relates the surface concentration $\phi_s \equiv \phi(z = 0)$ to the normal gradient of the bulk profile:

$$\kappa \frac{d\phi}{dz}\bigg|_{z=0} = \frac{\partial f_s}{\partial \phi_s} = -\mu_{\text{eff}}(C) + g\phi_s. \tag{S5}$$

Using the first integral of the bulk Euler–Lagrange equation [1, 2], the equilibrium profile satisfies

$$\frac{\kappa}{2}\left(\frac{d\phi}{dz}\right)^2 = W(\phi), \tag{S6}$$

where

$$W(\phi) \equiv f_b(\phi) - f_b(\phi_\infty) - \frac{\mu_\infty}{\nu_b}(\phi - \phi_\infty)$$

is the excess grand-potential density relative to the bulk reservoir state $\phi_\infty$ far away from the membrane surface, and $\mu_\infty$ is the chemical potential evaluated at $\phi_\infty$.

Evaluating this relation at the surface and combining it with Eq. (S5), we obtain the equilibrium conditions

$$\pm\sqrt{2\kappa W(\phi_s)} = -\mu_{\text{eff}}(C) + g\phi_s. \tag{S7}$$

Here, the sign of the gradient is fixed by the monotonicity of the equilibrium profile connecting surface concentration $\phi_s$ to the bulk reservior concentration $\phi_\infty$.

##### D. Mechanism: Curvature-mediated enhancement of surface prewetting

The formation of stable TFAM condensates on the membrane is governed by the interplay between bulk thermodynamics and the local surface potential. Our model reveals that membrane curvature functions not merely as a static geometric boundary, but as an effective thermodynamic field that locally modulates the phase diagram (see Fig. 4B in main text for details).

Mechanistically, the negative curvature ($H < 0$) characteristic of cardiolipin-enriched domains acts as a localized external field. Through the coupling term $\eta C\phi_s$, this geometric feature effectively shifts the local chemical potential $\mu_{\text{eff}}$, lowering the free energy barrier for the nucleation of the dense surface phase. Consequently, the prewetting transition—which would strictly require higher bulk concentrations on a flat membrane—is triggered locally within the curved domains at significantly lower bulk saturation levels (see Fig. 4B in main text for details). This curvature-induced "boost" explains the preferential emergence of thick condensate phases in concave pockets, while the the flat membrane remains in a thermodynamically distinct "thin" adsorption state (see Fig. 5D-E in main text for details).

#### Supplementary methods: Tomogram processing and curvature analysis

Additional implementation details for tomogram processing and membrane curvature analysis are provided below. Tilt series were preprocessed and manually inspected to remove low-quality frames prior to alignment in Warp 1.1.0 (`https://warpem.github.io`). Patch-tracking alignment was performed using AreTomo2 (`https://github.com/czimaginginstitute/AreTomo2`).

Missing-wedge correction and CTF deconvolution were applied using IsoNet v0.2 (`https://github.com/IsoNet-cryoET/IsoNet`). Automated membrane segmentation was performed using Membrain-seg (`https://github.com/teamtomo/membrain-seg`), followed by manual curation and membrane annotation in Amira 2022.1 (Thermo Fisher Scientific). Segmented volumes were converted to surface meshes using the Surface Morphometrics pipeline (`https://github.com/GrotjahnLab/surface_morphometrics`). Curvature metrics were computed using PyCurv with vector-voting-based curvature estimation. Intra- and inter-surface distances and surface orientation metrics were calculated using scripts provided in the Surface Morphometrics package. Quantitative curvature values were extracted from CSV format output files and used for downstream statistical analysis.

### Supplementary methods: Single-molecule tracking acquisition, code parameters and analysis

Time-lapse TIRF images of Texas Red-DHPE were acquired using a 561 nm laser at 2 mW with an exposure time of 10 ms at the interval at 12.5 ms. A region of $20 \times 20\,\mu$m was selected to minimize uneven illumination. After image preprocessing in Fiji, single-molecule trajectory analysis was performed using custom Python scripts adapted from the Octane single-particle tracking framework. The probability density histogram of step size was fitted to the following equation:

$$p(r) = \sum_{i=1}^{N} \alpha_i \frac{r}{2D_i\Delta t} \exp\left(-\frac{r^2}{4D_i\Delta t}\right), \tag{S8}$$

where N represents the number of populations (maximum of 2 for this study), $\alpha_i$ denotes the portion of the population, $D_i$ is the diffusion coefficient, and $\Delta t$ is the exposure time.

The MSD was calculated using the formula:

$$\mathrm{MSD}(t) = \frac{1}{N}\sum_{i=1}^{N} \left|\mathbf{x}_i(t) - \mathbf{x}_i(0)\right|^2, \tag{S9}$$

The MSD curve was fitted to an anomalous diffusion model:

The MSD curve was fitted using an anomalous diffusion model,

$$\mathrm{MSD}(\tau) = K_\alpha \, \tau^\alpha, \tag{S10}$$

where $K_\alpha$ represents the generated diffusion coefficient, which can be converted to the actual diffusion coefficient based on specific movement categories. Values of $\alpha$ close to 1 indicate free diffusion, $\alpha < 1$ suggests confined movement, and $\alpha > 1$ indicates directed movement. Given the complexity of heterogeneous movements, the study of lipid and membrane properties were classified by MSD, step size distribution, and track classification.

#### Supplementary methods: AFM characterization and mechanical analysis

Topographic imaging of membrane before and after Tfam titration was performed using an AFM (MFP-3D, Asylum Research) operated in AC water topography mode using a silicon nitride AFM probe. Cantilevers were manually tuned before measurement. Typical drive frequency was $\sim 5.8$ kHz, drive amplitude 3 V and sweep width 2 kHz. Images were acquired over $20\,\mu\text{m} \times 20\,\mu\text{m}$ with $512 \times 512$ pixels at 0.5 Hz scan rate. The setpoint was $\sim 45\%$ of free amplitude and optimized to minimize ghosting and tip-induced perturbation.

Mechanical properties of Tfam surface condensates were measured in contact mode. Cantilevers were calibrated by the Sader method. Force maps were acquired with a 200 pN trigger force and $0.1\,\mu$m ramp distance (AR SPM v16). A $40 \times 40$ grid over $5\,\mu\text{m} \times 5\,\mu\text{m}$ was measured at $0.125\,\mu$m spacing. Force mapping was conducted at room temperature with a 2 Hz rate and engagement velocity of $0.4\,\mu$m/s. Young's modulus was obtained using a Hertz model fit ($20-100$ pN fitting range on retraction). Distributions were analyzed and found to be consistent with Gaussian behavior with condition-dependent means and variances.

#### Key Resources

All resources applied in the experimental session are indicated in the table.

Table 1: Supplementary Table 1 — Key Resources Table.

| REAGENT or RESOURCE | SOURSE | IDENTIFIER |
|---|---|---|
| Coverslip | Marienfeld | Round: 0117650; Square: 0107052 |
| Ultrasonics Processor | XIAO MEI CHAO SHENG | 650T, 6mm |
| UV cleaner | Shanghai Zhongbin Technology CO., LTD | CCI UV-250MC |
| All lipids | Avanti | — |
| DNA | GENEWIZ INC. | — |
| Ultra-pure water machine | Hyperpurex | XUE-10/20UF |
| Albumin from Bovine Serum | Aladdin | A116563 |
| NeutrAvidin Protein, DyLight 405 | Invitrogen | 22831 |
| Ni-NTA beads 6FF | Smart-lifesciences | SA005005 |
| Size exclusion chromatography | Cytiva | Superdex 200 Increase 10/300 GL |
| Heparin-Sefinose column | Sangon Biotech | C600931-0005 |
| Alexa Fluor NHS Ester | Invitrogen | 488: A20000; 680: A20008 |
| Micro Bio-Spin Columns with Bio-Gel P-6 | Bio-Rad | — |
| Nanodrop | Thermo Fisher Scientific | — |
| SH-SY5Y | A gift from Junying YUAN's Lab | — |
| DMEM basic (1x) | Gibco | 6124314 |
| 200 mM Glutamine solution | Agilent | 103579-100 |
| Trypsin-EDTA solution | Biosharp | BL512A |
| DPBS | Biosharp | BL310A |

| REAGENT or RESOURCE | SOURCE | IDENTIFIER |
|---|---|---|
| Lipofectamine 2000 | Invitrogen | 11668019 |
| Polyethylenimine | YEASEN | 40815ES03 |
| G418 | Beyotime | B528413-0152 |
| Puromycin | Sangon Biotech | E607054 |
| MPP+ | Innochem | (CAS) 36913-39-0 |
| PK Mito Orange, PK Mito DeepRed | Genvivo | PKMO-2, PKMDR-1 |
| pET-28a, pcDNA3.1(+), lenti-CRISPRv2, psPAX2, pMD.2G | GENEWIZ INC. | — |
| Cell Mitochondria Isolation Kit | Beyotime | C3601 |
| Q5, T4-Ligase, Not1, Xho1, BamH1, BsmBI-v2 | NEB | — |
| Taq | Sangon Biotech | — |
| Opa1 antibody | BD Bioscience | 612606, 612607 |
| Tfam antibody | Abmart | PK05078 |
| Actin antibody | Abmart | P30002 |
| Beta-Tubulin antibody | Thermo Fisher | MA5-16308-HRP |
| TIRF module | FluoCa | — |
| Visview software | Olympus | — |
| Fluorescence Correlation Spectroscopy module | LightEdge Technologies Ltd. | — |
| EMCCD camera (TIRF) | Oxford Instrument | — |
| HIS-SIM microscopy | CSR biotech | — |
| Finer Imaging processing | CSR biotech | — |
| Confocal microscopy (FV4000) | Evident | — |
| STED and STED FLIM | Abberior Facility Line | — |
| AFM | Asylum Research | — |
| AFM probe | Budget Sensors | SiNi, 0.27 N/m |
| SICM | Eaglenos Sciences, Inc | — |

Table 2: Supplementary Table 2 — Oligonucleotide and sgRNA sequences.

| NAME | SEQUENCE (5′ to 3′) | 5′ MODIFY | 3′ MODIFY |
|---|---|---|---|
| 40-FC | ACAACGATCCAGCTGTCACTGAACCTAGATGTAAGTCGTC | — | Cholesterol |
| 40-RB | GACGACTTACATCTAGGTTCAGTGACAGCTGGATCGTTGT | — | Biotin |
| 40-RBT | GACGACTTACATCTAGGTTCAGTGACAGCTGGATCGTTGT | Texas Red | Biotin |
| 80-FC | CGTTGATGCGAGCTAGGTCCGATGCTACGTTCATGGTCACCTCGAGCGAAGTCACTTACGGTCAGTCAGGTATCCGTACG | — | Cholesterol |
| 80-RB | CGTACGGATACCTGACTGACCGTAAGTGACTTCGCTCGAGGTGACCATGAACGTAGCATCGGACCTAGCTCGCATCAACG | — | Biotin |
| 80-RBT | CGTACGGATACCTGACTGACCGTAAGTGACTTCGCTCGAGGTGACCATGAACGTAGCATCGGACCTAGCTCGCATCAACG | Texas Red | Biotin |
| ND1-F (in-vitro) | GGCTATATACAACTACGCAA | — | Alexa Fluor 680 |
| ND1-R (in-vitro) | AGGTGGCTAGAATAAATAGG | — | — |
| ATPase6-F (in-vitro) | CAAACTACCACCTACCTCCCTC | Alexa Fluor 647 | — |
| ATPase6-R (in-vitro) | TCCCGTATCGAAGGCC | — | — |
| RPLPO-F (qPCR) | GGAATGTGGGCTTTGTGTTC | — | — |
| RPLPO-R (qPCR) | CCCAATTGTCCCCTTACCTT | — | — |
| ND1-F (qPCR) | ATGGCCAACCTCCTACTCCT | — | — |
| ND1-R (qPCR) | CTACAACGTTGGGGCCTTT | — | — |
| MT-F (qPCR) | CACCCAAGAACAGGGTTTGT | — | — |
| MT-R (qPCR) | TGGCCATGGGTATGTTGTTAA | — | — |
| sgRNA-F | CACCGTCGGGAATTTGATCTCACCA | — | — |
| sgRNA-R | AAACTGGTGAGATCAAATTCCCGAC | — | — |

### Extended Data

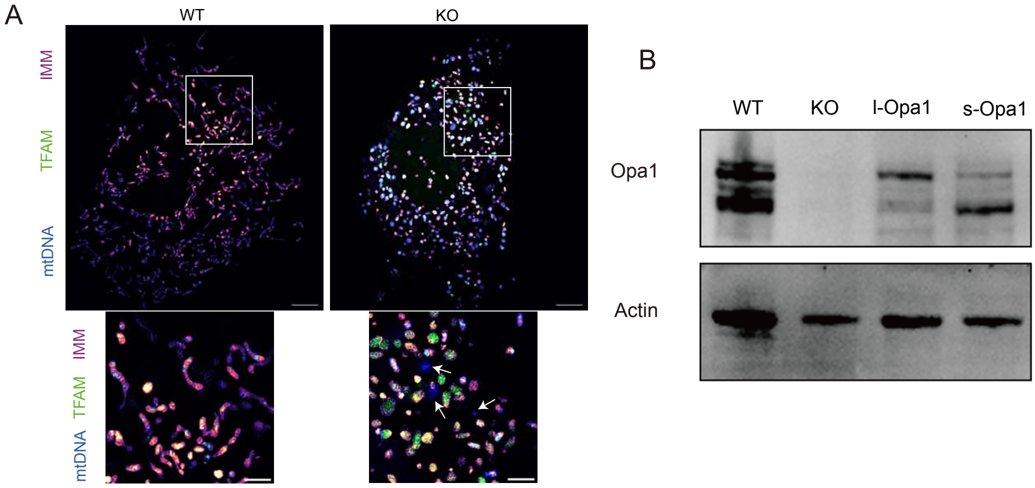


**Extended Data** 1: **IMM structure modified by Opa1 is associated with mtDNA releasing. A** Live cell SIM images of cells labeled for Tfam (green), mtDNA (Cyan) and the IMM (magenta)shows that in WT cells discrete Tfam puncta localizes well with mtDNA signals within mitochondrial matrix (Extended figure 1A, left), Opa1 KO cells however, exhibits disorganized IMM, where Tfam signal fills in the whole mitochondrial matrix, with mtDNA signal observed in plasma (Extended figure 1A, right, illustrated as pointed signals in the enlarged image). Top row scale bars: 5$\mu$m; Middle row scale bars: 2$\mu$m. **B** Immunoblotting results of WT mTec cells, Opa1 ko cells as well as l- and s-Opa1 rescue.

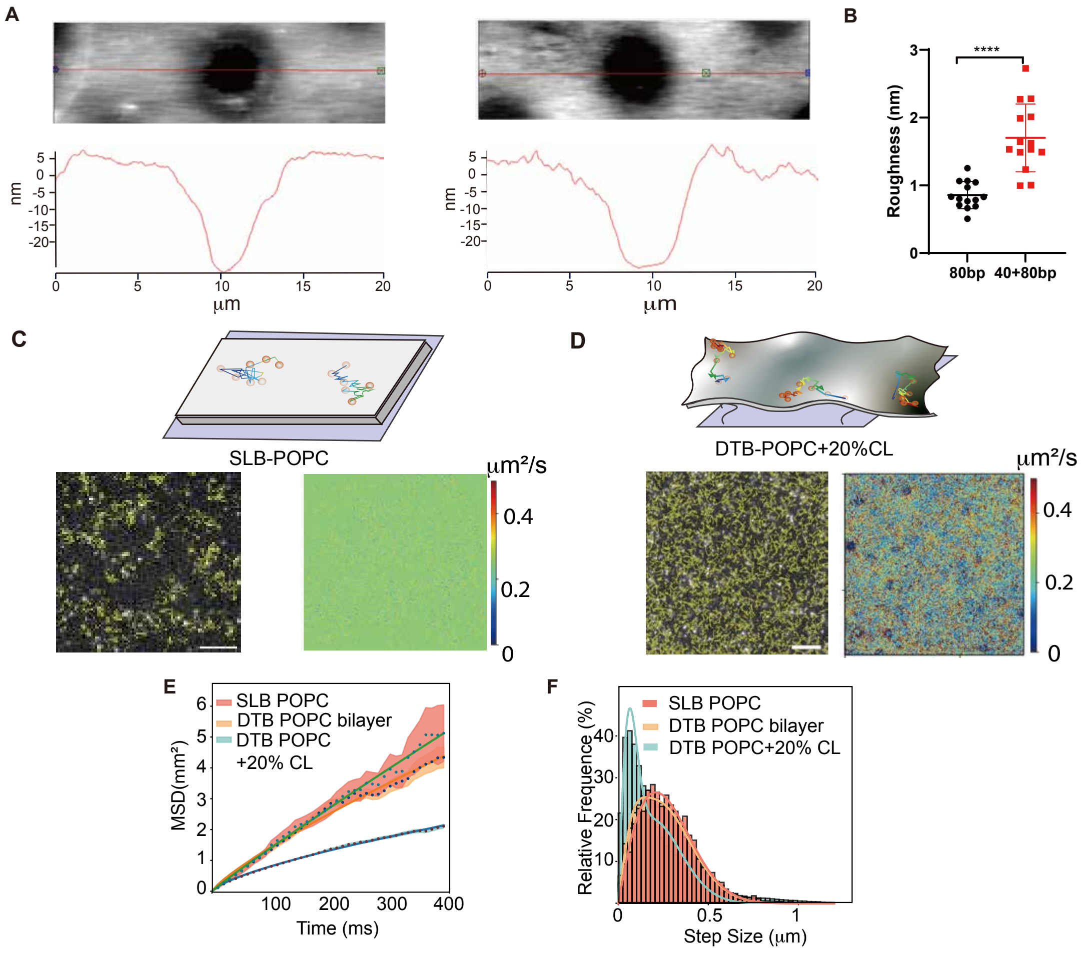


**Extended Data** 2: **Validation of DNA tethering effect and biophyisical characterization of DTBs.** **A**AFM images of DTB at patterned surface with homogeneous 80BP DNA tether (left) and heterogenous 40+80BP DNA tethering. The height difference between untreated substrate (dark) and bilayer coated region(bright) showed that DNA tethering successfully lifts the bilayer from substration. In addition, by adjusting the length double strain, the local curvedness shows distinguishable differences **B**. Single molecular tracking results showed that although all molecules are accessible in both no tethered and tethered bilayer, diffusion of probe lipids on SLBs are relatively homogeneous**C**, whereas tethered bilayers (80BP only,**D** showing localized diffusion traps where local diffusion coefficient of the bilayer is relatively low. This results are confirm by statistic analysis of MSD and step size distribution **E** and **F**.

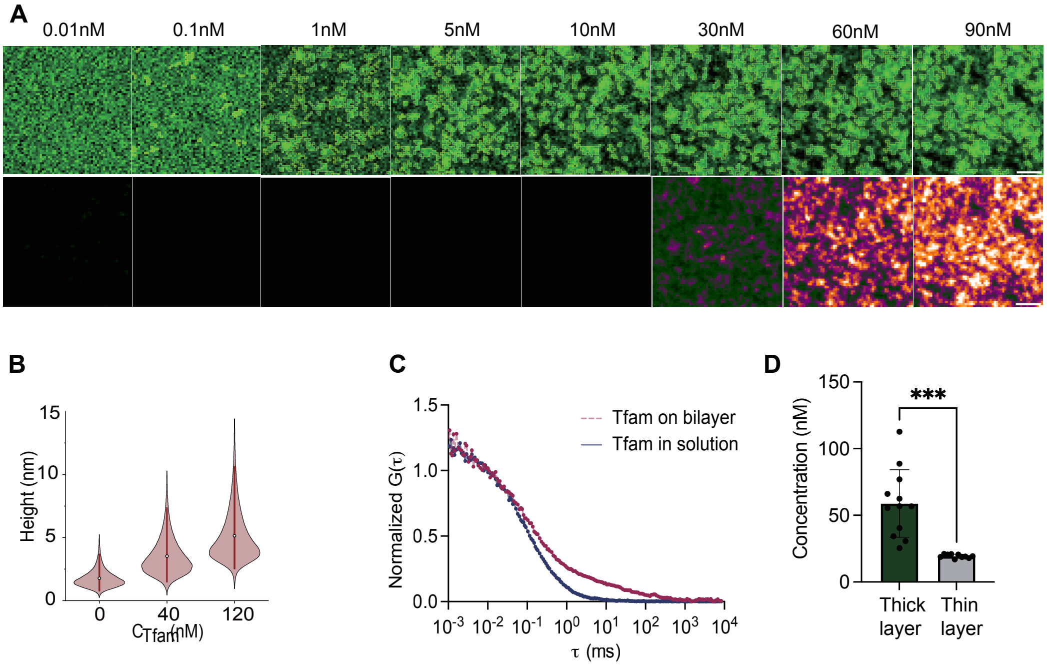


**Extended Data 3: Tfam titration on the surface of DTB. A** Raw fluorescent data obtained at the single localtion with Tfam titrated to the bilayer presented with original signal (upper) rwo and adjusted intensity signal (lower row). **B** Distribution of Z scale extension of bilayer and Tfam condensates during titration characterized by AFM analysis. **C**, normailized FCS autocorrelation curved of Tfam on bilayer and in solution, indicating a significant slower diffusion of Tfam at the surface of bialyer.**D** Tfam concentration measured by FCS of thin/thick with bulk concentration of 80nM,indicating the distinguishable protein concentration echos fluorescent signal obtained by TIRF.

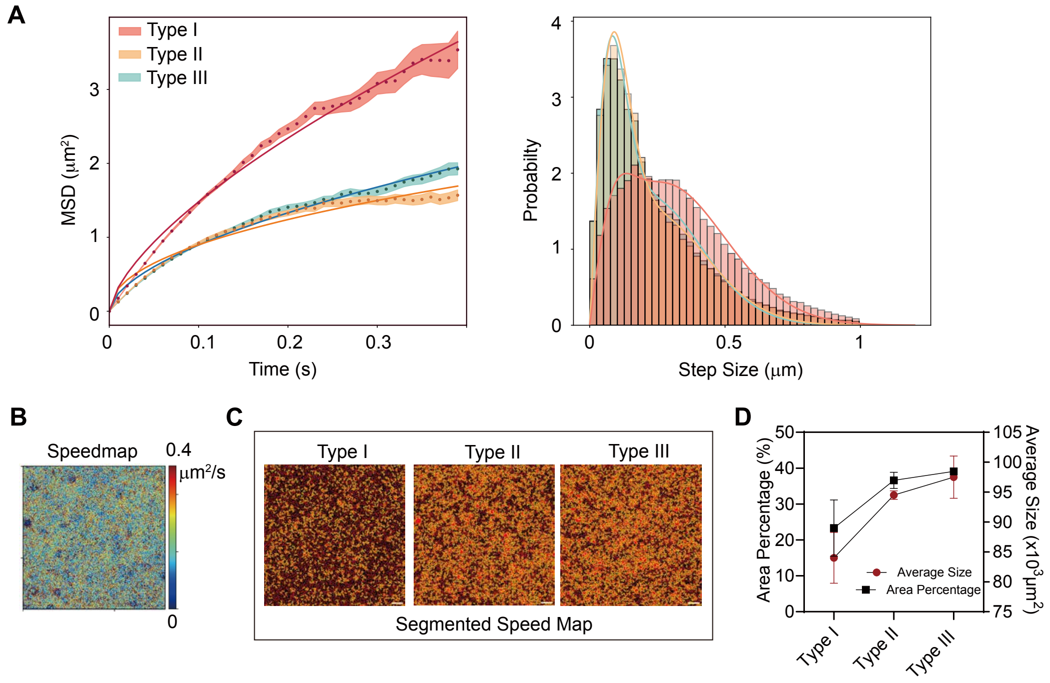


**Extended Data** 4: **Biophysical characterization of Type I, II, III bilayers labeled with fluorescence label (A)** MSD and step size distribution of single molecular tracking experiments obtained from Type I, II and III bilayers showed significant obstructed diffusion are observed with elevated membrane curveness. **B** Examplary speed map obtained from bilayer. **C** segmentation of the speedmap with water threshold method showing low diffusion areas elevated with membrane curvature, which is quantified in **D**.

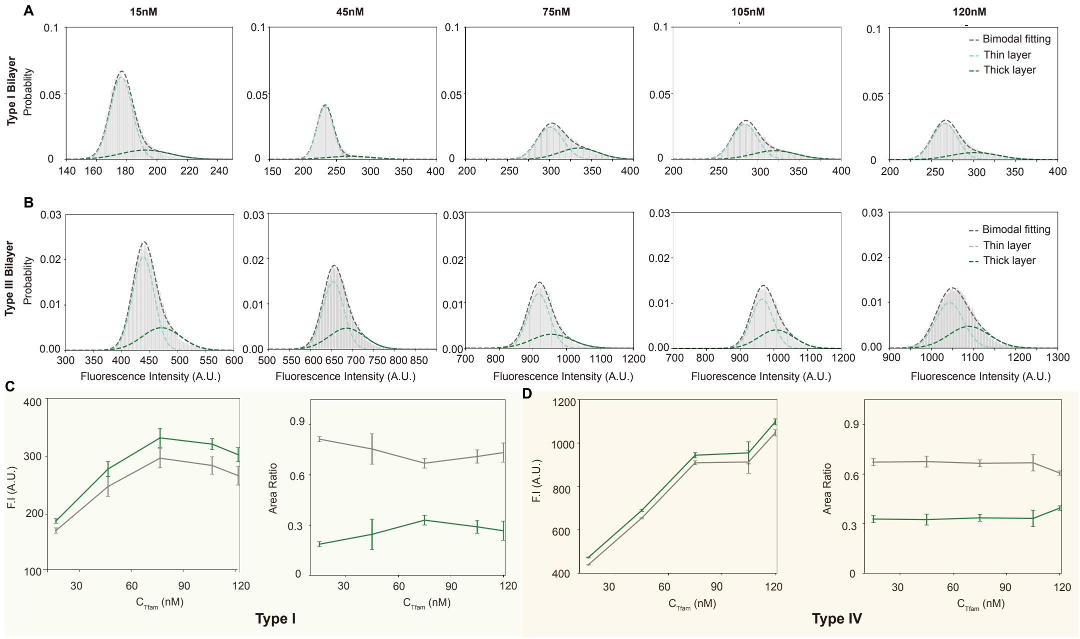


**Extended Data** 5: **Effects of membrane curvedness on Tfam condensation A** Exemplary bimodel fitting results of Tfam titrated on Type I and III bilayers **B** . The statistic results of Tfam accumulation (F.I) and area ratio at different bulk Tfam concentrations on Type I and Type IV bilayers are shown in **C** and **D**, respectively.

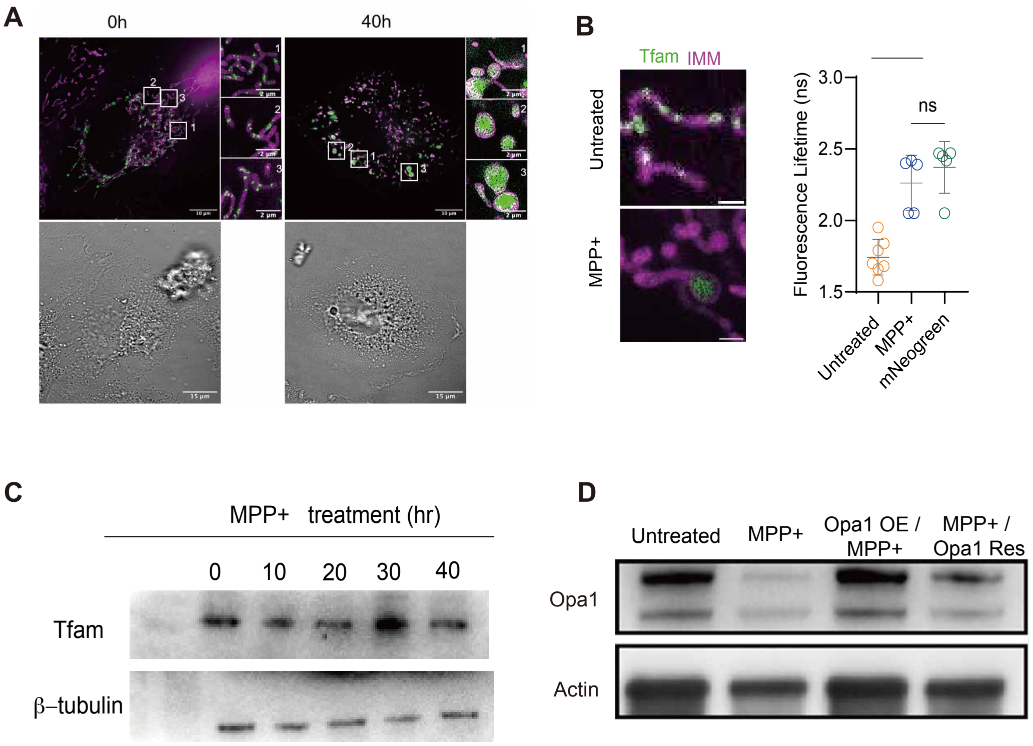


**Extended Data** 6: **A** SIM and DIC images of SH-SY5Y cells before and after 40 hours of MPP+ treatement. As Tfam shows a enlarged spreading area in mitochondria, STED-FLIM results of Tfam showed that Tfam-mNeoGreen are upacked from the condensed nucleoid as illustrated by elevated fluorescence lifetime that is identical to mNeogreen in solution. **C** Illustrate Tfam level stayed relatively constant upon MPP+ treatement, whereas Opa1 rescuing conditions were confirmed in **D**.

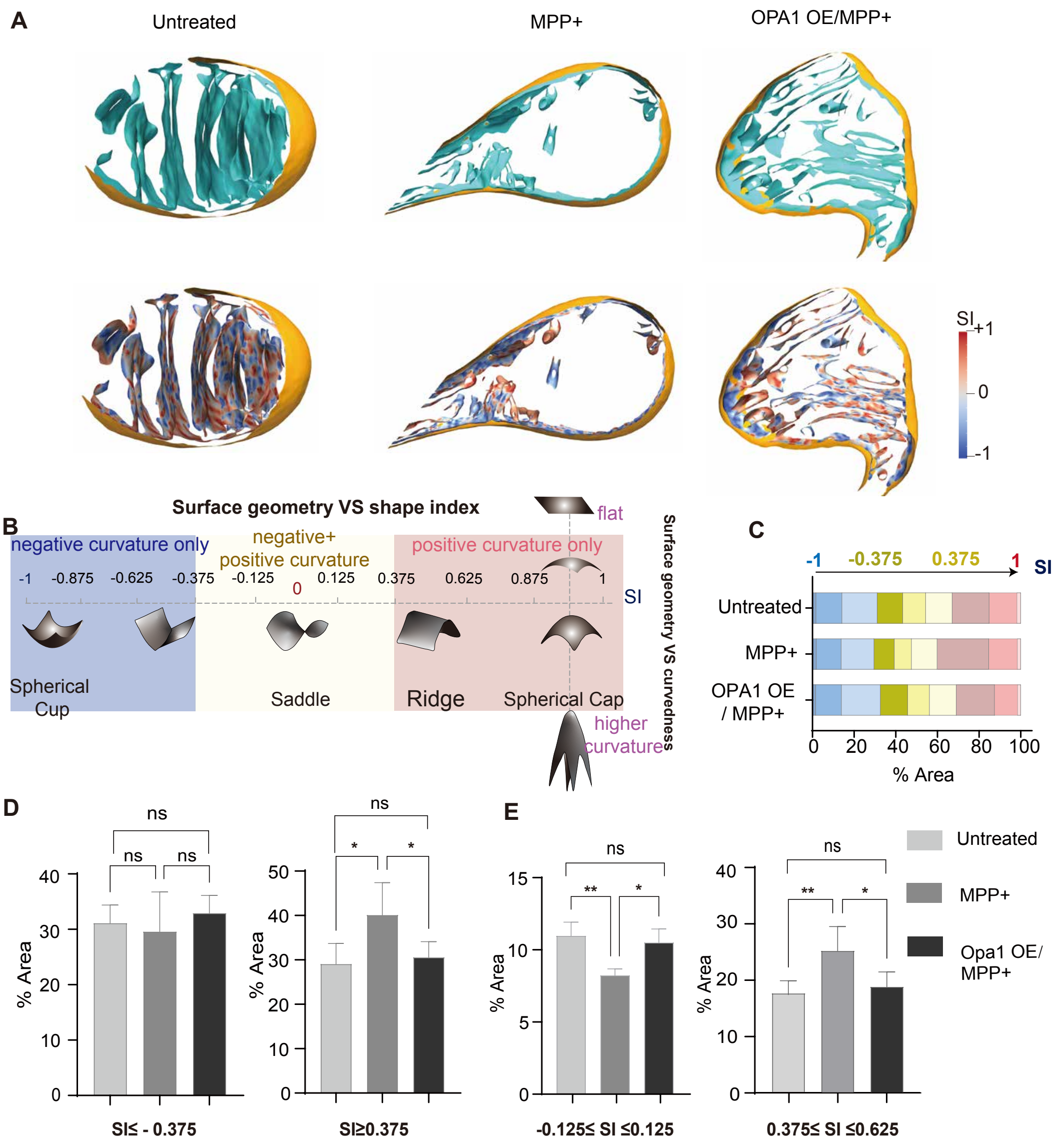


**Extended Data 7: IMM curvature shifts from neagtive to positive upon MPP+ treatement and partically recued by OPA1 OE A** 3D surface reconstructions of mitochondria under untreated, MPP+ (36 h), and Opa1 OE/MPP+ conditions. Top row: segmented membranes; Bottom row: corresponding Shape Index (SI) maps, color-coded by local morphology. **B** Shape Index reference key illustrating the classification of membrane curvature. SI = -1 represents a spherical cup (pure negative curvature); SI between -0.375 and 0.375 represents a saddle shape (mixed negative and positive curvature); and SI $>$ 0.375 to 1 represents a spherical cap (pure positive curvature).**C** Quantitative distribution of membrane area across SI values of each treatment condition, displayed as percentage of total IMM area. **D**.Quantification of IMM area fractions specifically at the transition geometries: (i) Saddle region (mixed curvature) and (ii) Ridge region (positive curvature, **E**)
